# Accuracy in near-perfect virus phylogenies

**DOI:** 10.1101/2021.05.06.442951

**Authors:** Joel O. Wertheim, Mike Steel, Michael J. Sanderson

**Affiliations:** Department of Medicine, University of California San Diego, La Jolla, CA 92093, USA; Biomathematics Research Center, School of Mathematics and Statistics, University of Canterbury, Christchurch, 8041, New Zealand; Department of Ecology and Evolutionary Biology, University of Arizona, Tucson, AZ 85721 USA

**Keywords:** Perfect phylogeny, Homoplasy, Yule-Harding model, virus, SARS-CoV-2, Ebola virus, Zika virus

## Abstract

Phylogenetic trees from real-world data often include short edges with very few substitutions per site, which can lead to partially resolved trees and poor accuracy. Theory indicates that the number of sites needed to accurately reconstruct a fully resolved tree grows at a rate proportional to the inverse square of the length of the shortest edge. However, when inferred trees are partially resolved due to short edges, “accuracy” should be defined as the rate of discovering false splits (clades on a rooted tree) relative to the actual number found. Thus, accuracy can be high even if short edges are common. Specifically, in a “near-perfect” parameter space in which trees are large, the tree length *ξ* (the sum of all edge lengths), is small, and rate variation is minimal, the expected false positive rate is less than *ξ*/3; the exact value depends on tree shape and sequence length. This expected false positive rate is far below the false negative rate for small *ξ* and often well below 5% even when some assumptions are relaxed. We show this result analytically for maximum parsimony and explore its extension to maximum likelihood using theory and simulations. For hypothesis testing, we show that measures of split “support” that rely on bootstrap resampling consistently imply weaker support than that implied by the false positive rates in near-perfect trees. The near-perfect parameter space closely fits several empirical studies of human virus diversification during outbreaks and epidemics, including Ebolavirus, Zika virus, and SARS-CoV-2, reflecting low substitution rates relative to high transmission/sampling rates in these viruses.

## Introduction

A “perfect phylogeny” is an evolutionary tree constructed from discrete character data in which no character state evolves more than once (Gusfield, 1997; Fernandez-Baca and Lagergren, 2003)—that is, homoplasy (Wake et al., 2011) is absent. Perfect phylogenies rarely exist for real-world datasets, but algorithms can be modified to search efficiently for “near-perfect” trees when a small amount of homoplasy is allowed (Fernandez-Baca and Lagergren, 2003; Awasthi et al., 2012). In this paper, we address how best to measure accuracy in such “near-perfect” trees, what factors guarantee accuracy is high, and whether real datasets with such minimal levels of homoplasy even exist.

The concept of perfect and near-perfect phylogenies played a key role in early attempts to understand the connections among phylogenetic tree reconstruction methods, such as maximum likelihood (ML), maximum parsimony (MP), and maximum compatibility. In a landmark paper, Felsenstein (Felsenstein, 1973) showed that a sufficient condition for ML and MP to infer the same tree was for the expected number of substitutions on edges of the tree to be very small. Then, “[i]f our assumption were true that evolutionary change is improbable during the relevant period of time, most characters should be uniform over the group. A few would show a single change of state during the evolution of the group. But only very rarely would we find more than one change of state, so that few or no characters would show convergence.” This last statement may have been the first hint of a probabilistic description of “near-perfect phylogeny”. This condition can be stated more formally as *ξ* ⩽ 1, where *ξ* is the expected number of substitutions per site summed over the entire tree (i.e., the tree length per site). Homoplasy is rare but has a non-zero probability of occurring.

Felsenstein’s concluding comment on near-perfect phylogenies was skeptical: “Real data is certainly not like this…” (Felsenstein, 1973). Homoplasy has since been viewed as a commonplace feature of phylogenetic datasets (Wake et al., 2011) and, reasonably enough, most phylogenetic theory has been developed with this sentiment as an implicit assumption. However, extensive surveys of genetic diversity in RNA viruses have revealed that some viral phylogenies, particularly those associated with outbreaks and epidemics, do exhibit small per site total tree lengths consistent with near-perfect phylogenies (Dudas and Bedford, 2019). These datasets often comprise full-length viral genomes from RNA viruses, which are typically 10–30 kb in length and have a substitution rate of around 10^−3^ substitutions/site/year.

The potential of these data to yield fully resolved phylogenies has been of particular interest in epidemiology, because internal nodes in viral trees represent transmission events (Campbell et al., 2018; Grubaugh et al., 2019; Dudas and Bedford, 2019). This motivates placing a premium on minimizing false negatives (i.e., on deciphering all such transmission events). However, understanding the false positive rate remains a key issue in characterizing phylogenetic accuracy overall (Felsenstein and Kishino, 1993).

Here we explore what assumptions comprise “near-perfect” phylogenies and decouple the false-positive and false-negative components of accuracy in such trees. In particular, by focusing on a mathematically tractable case in which tree size is large yet tree length is small, we will show that the false positive rate can be very good, even when the false negative rate is not: most of the clades inferred are probably correct, even though the tree may be only partly resolved. We also survey a set of viral phylogenies that have many properties of this near-perfect space and estimate their accuracy. Finally, we briefly consider phylogenetic “support” measures in relation to accuracy in near-perfect data. Whereas accuracy relates to the overall performance of a tree estimator relative to the true tree, support relates to the probability of making a mistake in deciding about some aspect of that tree—typically the presence of a particular split—using a statistically based decision rule such as the bootstrap support value or a posterior probability (Felsenstein, 1985; Felsenstein and Kishino, 1993; Hillis and Bull, 1993; Efron et al., 1996; Susko, 2008, 2009; Alfaro and Holder, 2006; Simmons and Norton, 2014).

This paper is organized as follows. “Materials and Methods” are divided into two parts: first, mathematical theory (with proofs in the Supplement), and second, simulation protocols, data, and data analysis. “Results” begin with a more expository description of the theory, illustrated with simulation results, and then describes results from analyses of robustness and support, and data analyses. Following these is the Discussion.

## Materials and Methods I. Theory

### Definitions of Accuracy

Given a true unrooted binary tree, *T*, and an estimated tree, 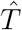, a strict measure of accuracy is just 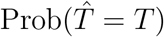 (Huelsenbeck and Hillis, 1993; Erdös et al., 1999). In large trees it is useful to measure partial agreement, such as the proportion of nontrivial splits on 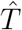 that are also on *T*, out of a possible *n* – 3 (Yang, 1998).

A still more nuanced definition of accuracy is useful when either *T* or 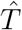 is only partially resolved (not binary), that is, when the number of nontrivial splits, *C*(*T*), is less than *n* – 3 (Warnow, 2013). Let *N_FP_* be the number of splits on 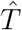 but not *T* (false positives), and let *N_FN_* be the number of splits on *T* but not 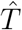 (false negatives). When both trees are binary, *N_FP_* = *N_FN_* (Berry and Gascuel, 1996; Smirnov and Warnow, 2021); otherwise they can contribute differentially to error. The Robinson–Foulds (RF) distance (Robinson and Foulds, 1981), *d_RF_* = *N_FP_* + *N_FN_*, combines both errors in one measure of overall accuracy. Here we distinguish between these errors explicitly by defining false positive and negative rates (Smirnov and Warnow, 2021):

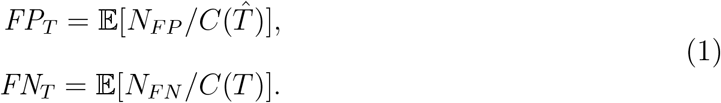

Both error rates are expectations over some generating model for the data, described next.

### Evolutionary Model

Let *B*(*n*) denote the set of unrooted binary phylogenetic trees with leaf set [*n*] = {1, 2,…, *n*}. Note that a tree *T* ∈ *B*(*n*) has 2*n* – 3 edges. Consider a Jukes-Cantor model (JC69; Felsenstein, 2004), with rate parameter λ, in which the probability of a state change between the endpoints of an edge *e*, denoted *p_e_*, is given by *p_e_* = *p*, where 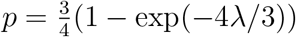. Assume further that all edges have the same value of λ. Let *ξ* denote the expected number of state changes per character in *T*. Thus *ξ* = λ · (2*n* – 3).

A *character* refers to the assignment of states to the taxa at a given site of an alignment. We will say that a character evolves ‘perfectly’ on *T* if there is a single change of state across one interior edge (say *e*) and no change of state on any other edge of *T*. Thus, a character that evolves perfectly on *T* is homoplasy-free, and the two notions are equivalent for binary characters. However, for multi-state characters, the notion of a perfectly evolved characters is stronger than that of being being merely homoplasy-free. We deal here with this stronger notion for two reasons: firstly, it simplifies the mathematical analysis, and second, the expected proportion of homoplasy-free characters that not perfectly evolved under the models we consider tends to zero as the number of taxa becomes large.

We will say that a character *f* evolves on *T* with *c edge changes on e*_1_, …, *e_c_* if state changes occur on edges *e*_1_, …, *e_c_* and on no other edge of *T*. More briefly, we say that *f* evolves on *T* with *c* edge changes if *f* evolves with *c* edge changes for some set of *c* distinct edges of *T* (mostly we will deal with the case *c* =2).

Recall that a *split* refers to a bipartition of the leaf set [*n*] into two nonempty subsets (and splits are induced by binary characters). A character that has evolved perfectly on *T* produces a split, and these splits (across a set of perfectly evolved characters) are compatible and so form a (generally unresolved/non-binary) tree on leaf set [*n*].

### Probability of False Splits

Suppose that *m* characters evolve on *T* and that, of these *m* characters, *k* of them are perfectly evolved on *T* (note that more than one of these characters may correspond to the same split of *T*). Next, consider a single additional character *f* which has evolved on *T* with 2 edge changes, on *e*_1_, *e*_2_ (there is no restriction that these must be interior edges). Under certain conditions, the MP tree for these characters will include a false split (false positive)—a split not on *T* (Fig. 1). In particular, a false split occurs if no perfect character changes state along the path between *e*_1_ and *e*_2_ (see Lemma 1 in the Supplementary Information).

**Fig. 1.**
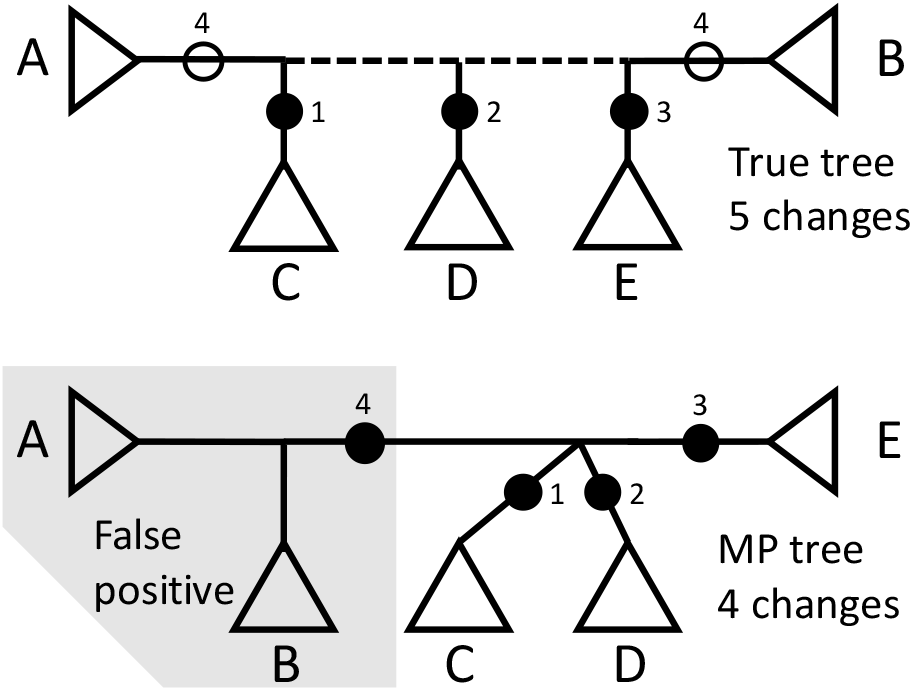
How a false positive split is inferred by maximum parsimony (MP). On true tree (top) sites 1–3 are binary and “perfect”; that is, they have only a single change (locations marked by black circles), but site 4 is binary and homoplastic, changing twice (open circles). The dotted line is the path between the two homoplastic changes in site 4. As long as perfect sites do not change along the dotted line path, a false positive split is inferred on the MP tree (bottom).

Let 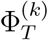 be the probability that a character *f* that has evolved on *T* with 2 edge changes generates a false split under MP, which means:

**(C-i)** it is a binary character,
**(C-ii)** the corresponding split is not a split of *T*, and
**(C-iii)** the split described by *f* is compatible with *k* characters that are perfectly evolved on *T* (by the Markovian process described above).

Given a tree *T* ∈ *B*(*n*), let *d_T_*(*e*_1_, *e*_2_) denote the number of edges of *T* that lie strictly within the path between *e*_1_ and *e*_2_ (i.e., excluding *e*_1_ and *e*_2_). Thus, *e*_1_ and *e*_2_ are adjacent if and only if *d_T_*(*e*_1_, *e*_2_) = 0. In addition, let *φ_T_* = (*φ_T_*(0), *φ_T_*(1), …, *φ_T_*(*n* – 3)), where *φ_T_*(*i*) is the number of (unordered) pairs of edges {*e*, *e*′} of *T* for which *d_T_*(*e*, *e*′) = *i*. Finally, for *i* between 1 and *n* – 3, let

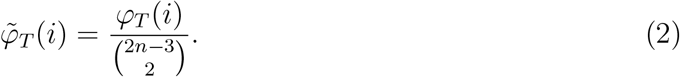

The probability of a false split is then given by the following theorem (see SI for proof).

#### Theorem 1

For each *T* ∈ *B*(*n*), and *k* ⩾ 1 we have:

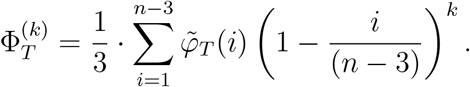

Theorem 1 shows that for fixed *k* and *n*, the shape of *T* plays a significant role in determining 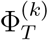 in particular, unbalanced trees (such as caterpillars) will have a smaller value of 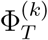 than more balanced trees. Indeed, it is possible to calculate the value of 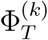 exactly for the two extreme cases of caterpillar trees and fully-balanced trees to determine the extent of this dependence (see SI).

### Estimating the Expected False Positive Rate

Given a binary phylogenetic tree *T*, and *m* characters evolved randomly on *T* by the model described earlier, the *false positive rate* (*FP_T_*) is the expected value of the ratio of false splits to all splits in the estimated tree (Eqn. 1; here we assume that if the reconstructed tree is a star, this proportion [which is technically 0/0] is zero). Recall that *ξ* is the expected number of state changes in the tree *T* per character, under the model described earlier. *FP_T_* is a function of the three parameters *T* (specifically, its shape and number of leaves), *m*, and λ (equivalently, *FP_T_* is a function of *T*, *m*, and *ξ*).

In general, it is mathematically complicated to describe *FP_T_* in terms of these parameters. However, when the number of leaves in a tree grows faster than the number of perfectly compatible characters, it is possible to state a limit result to provide an approximation to *FP_T_* for large trees.

In the following theorem, we consider the following setting:

I. *mξ* = Θ(*n^β^*) for some 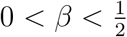, and
II. *mξ*^2^ = *O*(1),

where *O*(1) refers to dependence on *n* (thus *mξ*^2^ is not growing with *n*). Note that Condition (I) implies that the number of perfectly evolved characters grows with the number of leaves, but at a rate that is slower than linearly. Conditions (I) and (II) imply that *ξ* decreases as *n* increases.

In this setting, we show that the false positive rate is (asymptotically) of the form 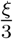 times a function Ω that involves *T* (via its shape), *m*, and *ξ*. If we now treat *ξ* as a variable, then for *ξ* = 0, the function Ω is close to 1 (for large *n*) and so *FP_T_* initially grows like *ξ*/3. However, as *ξ* increases, Ω begins to decline at an increasing rate, resulting in the false positive rate reaching a maximum value before starting to decrease.

To describe this result, we need to define this function Ω. Let

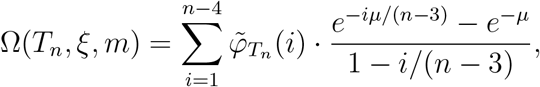

where:

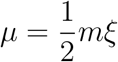

and where 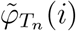 is given in Eqn. (2). For example, for any caterpillar tree, we have 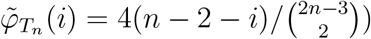.

Notice that Ω(*T_n_*, *ξ*, *m*) depends on *T_n_* only via the coefficients 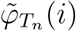, and this dependence is linear. Thus, if 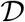 is a distribution on trees (e.g. the PDA or YH), then the expected value of Ω(*T_n_*, *ξ*, *m*) is given by:

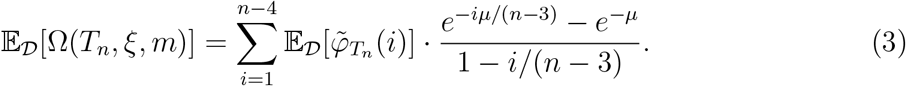

For the PDA distribution, the term 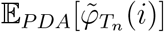 has an explicit exact value, namely,

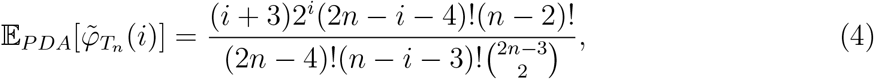

for all *i* between 1 and *n* – 3 (see SI for proof).

#### Theorem 2

For each *n* ⩾ 1, let *T_n_* be a binary phylogenetic tree with *n* leaves, and suppose that Conditions (I) and (II) hold.

i. 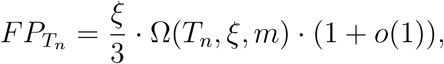

where *o*(1) is a term that tends to 0 as *n* grows.
ii. If *T_n_* is sampled from a distribution 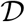 (e.g. PDA, YH), then the expected value of *FP_T_n__*, denoted 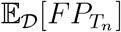, satisfies

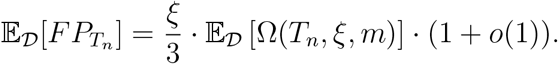

#### Remarks

Note that *FP_T_n__* depends only on the shape of the tree *T_n_* (and not on how its leaves are labelled), thus for a tree distribution 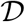 on either the class of caterpillar trees, or symmetric trees, we have 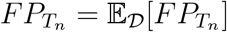.

Notice also from Fig. 3 that as *ξ* increases from 0 the estimate of 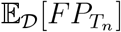 given by 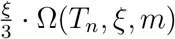 for the YH, PDA distributions and for symmetric trees initially increases (approximately linearly) with *ξ* but then begins to decrease with increasing *ξ*. By contrast, when *T_n_* has the caterpillar tree shape, the estimate of *FP_T_n__* appears to be constant as *ξ* increases from 0 (see Fig. 3). Indeed, when *T_n_* is a caterpillar tree, the expression for *FP_T_n__* in Theorem 2(i) reduces to the following remarkably simple expression as *n* becomes large:

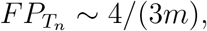

which is independent of *ξ* (and *n*). Details are provided in the SI.

## Materials and Methods II. Simulations, Data, and Data Analyses

### Main Simulation Pipeline

Simulations were run to assess goodness of fit and robustness of mathematical predictions under various regimes of model parameters and tree inference criteria (MP or ML), as well as to estimate expected accuracy in empirical data sets. Each of *R* simulation replicates (with *r* sub-replicate tree searches in each) consisted of the following sequence of steps: (i) generation of a random binary tree *T* with *n* leaves according to either a “proportional-to-distinguishable-arrangement” (PDA) or Yule-Harding (YH) model (Aldous, 2001) (as well as the two extreme cases of completely unbalanced caterpillar trees, and completely balanced symmetric trees); (ii) assignment of edge lengths of *T* according to a gamma distribution with shape parameter *α_e_* and mean 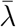; (iii) generation of a sequence alignment of *m* sites using either JC69, HKY or GTR models (using Seq-Gen v. 1.3.4, with base frequencies, rate matrix parameters, invariant site parameter or gamma shape parameter set or estimated from empirical data); (iv) reconstruction of estimated tree 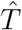 [using PAUP 4.0a (build 166) for MP with options ‘hsearch add=simple swap=no nreps=*r*;contree all/strict’; and using IQ-Tree2 (v. 2.0.6) (Minh et al., 2020) for ML with options ‘–m HKY+FQ -nt 1 -redo -mredo –polytomy -blmin 1e-9’, replicated *r* times, followed by strict consensus]; (v) tallying *N_FP_* and *N_FN_* from *T* and 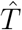 and computing error rates. Mean rates across replicates were then tallied. All steps except (iii) and (iv) used custom PERL scripts (available at https://github.com/sanderm53/pperfect). Generally, *R* was set to 1000 and *r* to 100.

### Support Simulations

Phylogenetic support measures were estimated in trees simulated via the main pipeline described above with *n* = 513, *m* = 1000, a JC69 model with no rate variation, and PDA random trees. Ten values of λ in the interval [10^−5^, 0.31622] were analyzed. PAUP was used for MP bootstrapping (same heuristic search as above but with 100 replicates × 10 subreplicates); IQTree2 was used (50 random tree replicates) for SH-aLRT (‘-alrt 1000’), aBayes, and ultrafast bootstrapping (‘-B 1000’), with additional options enforcing minimum branch lengths of 10^−9^ and collapsed polytomies. Mean support across replicates was computed.

Perfect four-taxon alignments were generated in which each of the five branches had a single, non-homoplastic nucleotide substitution in the alignment and all other sites were constant. Alignment lengths ranged between 40 nt and 30,000 nt. ML trees were inferred in IQTree2 with a JC69 model, minimum branch lengths of 10^−9^, and collapsed polytomies. Clade support was determined using ultrafast bootstrapping (10,000 replicates), SH-aLRT (10,000 replicates), and aBayes. Full Bayesian inference was also performed in MrBayes v3.2.7 (Ronquist and Huelsenbeck, 2003) with a single run per replicate of 2.5 million generations, with the first 10% of generations discarded as burnin.

Alignments for larger perfect symmetrical and asymmetrical (caterpillar) trees were generated with 8, 16, 32, 64, and 128 taxa. Each branch, including terminal branches, had a single nonhomoplastic nucleotide substitution in the alignment with all other sites constant. Alignment lengths ranged from 236 to 32,768 nt. ML trees were inferred as described above for the four-taxon alignments, and support was assessed by ultrafast bootstrapping, SH-aLRT, and aBayes.

### Virus Datasets

Viral phylogenies were obtained from the NextStrain (Hadfield et al., 2018) website (accessed 05 May 2020) (Table 1). Phylograms were downloaded for dengue virus, sengue virus serotype 1, Ebolavirus (Dudas et al., 2017), Enterovirus 68 (Dyrdak et al., 2019), measles morbillivirus, mumps virus, respiratory syncytial virus, West Nile virus (Hadfield et al., 2019), and Zika virus. In addition, we also analyzed an iatrogenic HIV-1 outbreak in Cambodia (Rouet et al., 2018) and the first wave of the SARS-CoV-2 epidemic in China (Pekar et al., 2021). The SARS-CoV-2 phylogeny is the ML tree used in Pekar (Pekar et al., 2021) (see Data S1 for list of GISAID Accession IDs). Publicly available genomic sequences (or genetic sequences for HIV-1) were downloaded from GenBank and aligned with mafft v7.407 (Katoh and Standley, 2013) (accession numbers can be found in Data S2).

**Table 1.** Parameters of 11 empirical virus phylogenies

| Abbreviation | Study | Leaves | Sites | Resolution |
| --- | --- | --- | --- | --- |
| DENV | Dengue virus | 1197 | 10264 | 0.8795 |
| DENV-1 | Dengue virus serotype 1 | 1067 | 10264 | 0.8160 |
| EBOV | Ebolavirus | 1610 | 18164 | 0.3632 |
| EV-D68 | Enterovirus 68 | 824 | 7293 | 0.8029 |
| HIV-1 | Human immunodeficiency virus type 1 | 189 | 1038 | 0.2193 |
| MeV | Measles morbillivirus | 109 | 15782 | 0.7009 |
| MuV | Mumps virus | 458 | 15154 | 0.2961 |
| RSV | Respiratory syncytial virus | 997 | 14986 | 0.6121 |
| SARS-CoV-2 | Severe acute respiratory syndrome coronavirus 2 | 583 | 29668 | 0.2324 |
| WNV | West Nile virus | 2512 | 10395 | 0.5960 |
| ZIKV | Zika virus | 543 | 10320 | 0.5453 |

False positive rates for the virus phylogenies were estimated with our simulation pipeline, setting parameters to values estimated from published trees and publicly available sequences used to construct them (Table 1, S1). IQTree2 was used to infer the six rate parameters of a GTR substitution model with empirical base frequencies, and either ASR variation following a gamma distribution with shape parameter 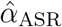, or an invariant sites model with parameter *p*_invar_ (‘GTR+F+G4’ or ‘GTR+F+I’). Model fit was assessed using the Bayesian Information Criterion (BIC) in IQTree2 (Kalyaanamoorthy et al., 2017).

Edge length (per site) variation was assumed to follow a gamma distribution: 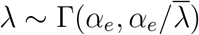 having mean 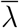 and variance 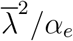. The distribution of substitutions is a mixture of Poisson and gamma distributions, which is a negative binomial with a variance to mean ratio of

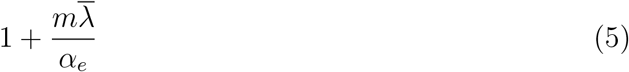

which was shown by Bedford and Hartl for an equivalent parameterization (Bedford and Hartl, 2008). Virus trees were preprocessed, setting any edge lengths < 1.1 × 10^−6^ to zero, assuming these reflected ML numeric artifacts. Then, 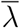 was estimated from the observed sum of per site edge lengths divided by 2*n* – 3, and Eqn. 5 was then used to estimate *α_e_*.

Ideally, we would fit the data to the random tree model, but standard methods either assume binary trees or model polytomies with an a priori assumption about the tree model itself (e.g., Bortolussi et al., 2006). Therefore, we repeated simulations using both PDA and YH models.

## Results

### Overview of Results on Accuracy

Simulations of tree inference with MP, over a large range of tree lengths, *ξ*, and other parameters illustrate several known results (Fig. 2) and perhaps a few less well known ones. First, resolution of the inferred tree increases with tree length. Second, “overall” accuracy, as measured by the RF distance, is optimal at an intermediate tree length, *ξ*\* (Yang, 1998; Bininda-Emonds et al., 2001; Steel and Leuenberger, 2017). Moreover, when *ξ* >> *ξ*\*, the false positive error rate, *FP_T_*, is similar to the false negative rate, *FN_T_*, as might be expected because the true and estimated trees are nearly binary; therefore *N_FP_* ≅ *N_FN_*.

**Fig. 2.**
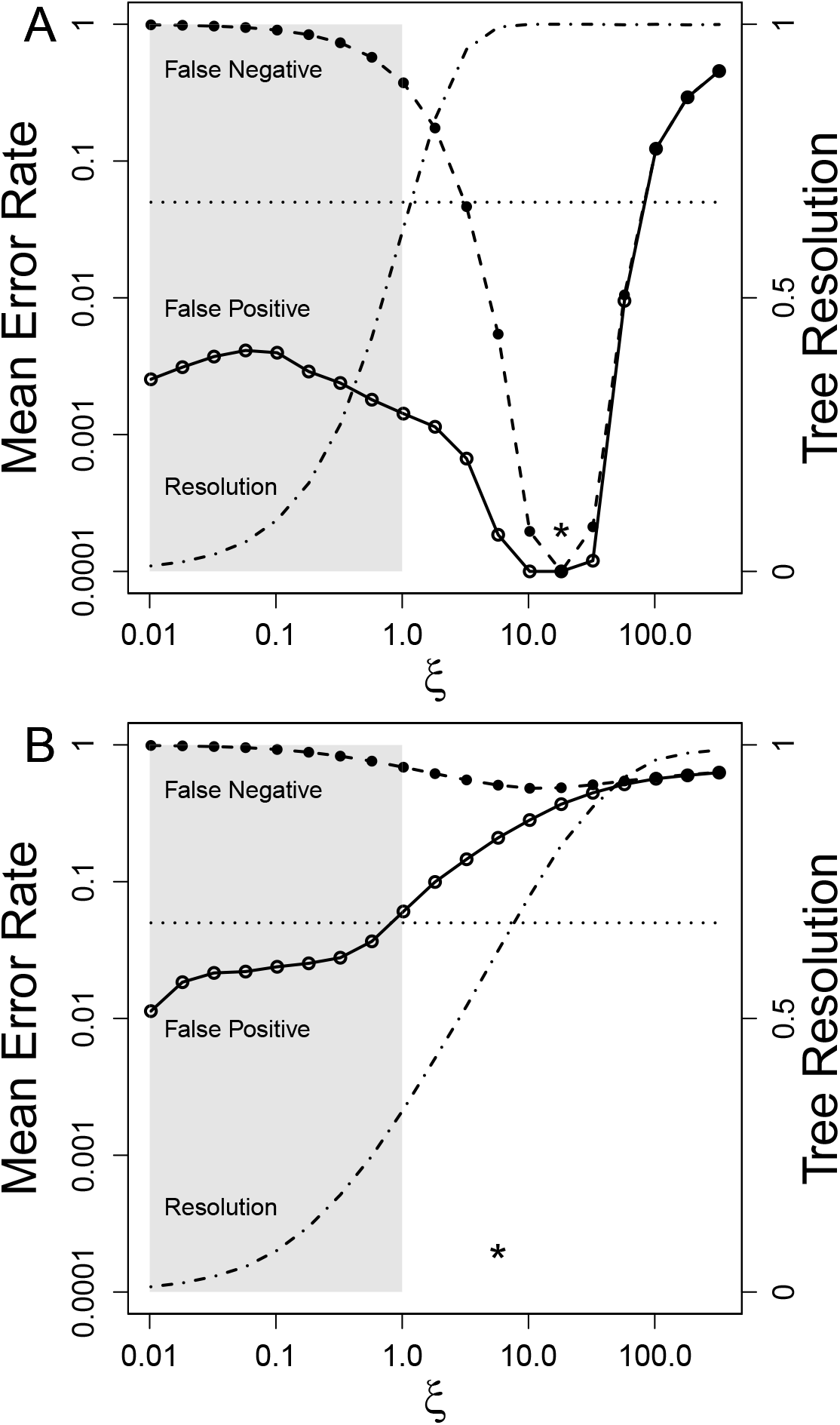
Accuracy of maximum parsimony phylogeny reconstruction in simulations over a wide range of tree length, *ξ*, and other parameters. Open circles are mean false positive error rates; closed circles are mean false negative error rates (log scale left); dashed sigmoidal curve is fractional resolution of estimated tree (linear scale right). Trees are generated by a random proportional-to-distinguishable-arrangement (PDA) model for 513 taxa, from which a sequence alignment length of 1000 sites is generated. The dotted horizontal line is placed at an error rate of 0.05. Asterisk marks the location of the optimal tree length with best overall Robinson–Foulds accuracy, *ξ*\*. Each point is mean of 1000 replicates × 100 sub-replicates (see Methods). “Near-perfect” values of *ξ* ⩽ 1.0 are shaded. A) JC69 model with no edge length or across-site-rate variation. Because of *y*-axis log scaling, two *y* values of zero were set to 0.0001. B) JC69 model with substantial edge length and across-site-rate variation, both modeled as a gamma distribution with shape parameters *α_e_* = *α_ASR_* = 0.25).

**Fig. 3.**
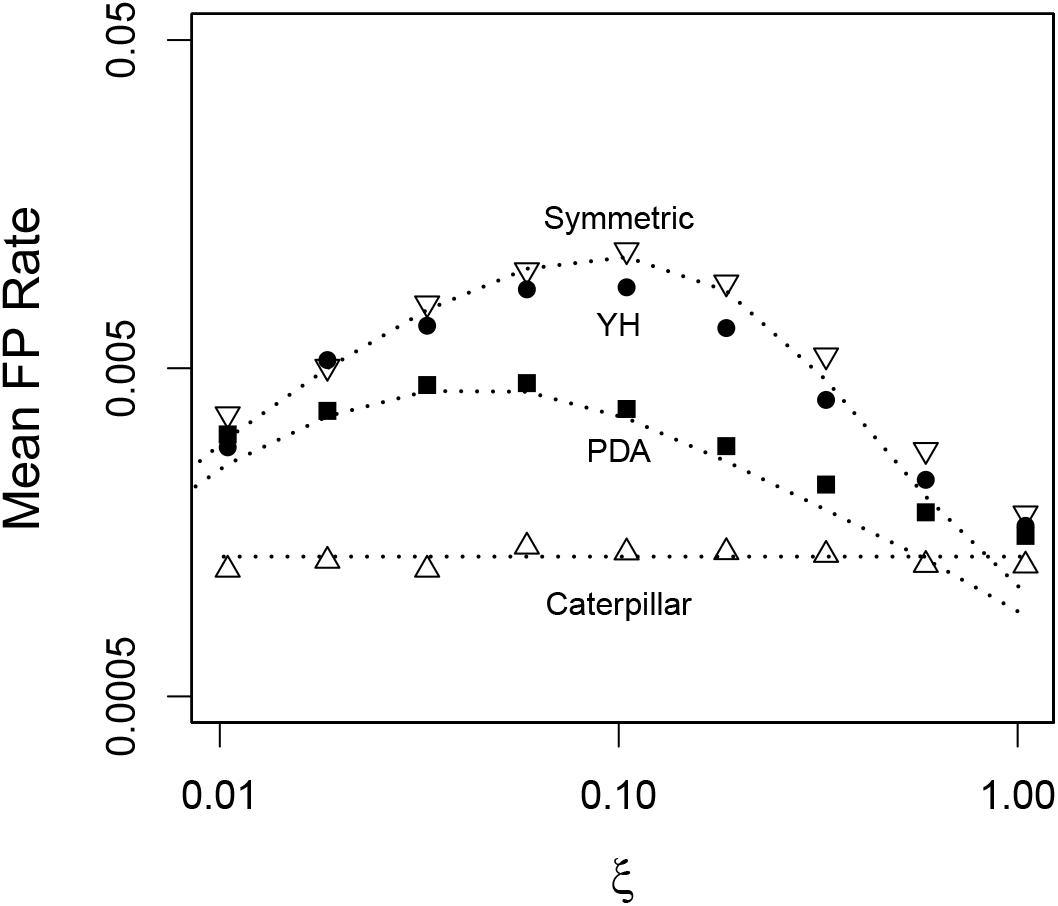
Mean false positive rate in four tree models. Fit to theoretical predictions from Equation 6 (or the limit expression of 4/3*m* for caterpillar trees: see Methods) are shown by dashed lines. Each point is mean of 1000 replicates × 100 sub-replicates. Simulation conditions were *n* = 513, *m* = 1000, with a JC69 model. Predicted values are not known for YH model.

However, when *ξ* << *ξ*\*, *FP_T_* << *FN_T_*, and the false positive error rate can remain quite good (< 0.05) over a large range of *ξ* even when the false negative error rate is very high. However, the range of tree lengths for which this result holds depends critically on rate variation across edges and sites. When *ξ* ⩽ 1, the false positive rate is low and insensitive to the presence of rate variation; but, when *ξ* > 1, the false positive rate is much more sensitive to rate variation—high when variation is present and low when absent (contrast Fig. 2A and Fig. 2B). In real-world data, as *ξ* increases, we expect that evidence of rate variation will become more apparent. Key elements of these findings can be shown analytically in a “near-perfect” zone described by a simple evolutionary model.

### Overview of the Mathematical Theory

First we define “near-perfect” more formally. Assume the data consist of an alignment of *m* independent and identically distributed nucleotide sites that have evolved according to a Jukes-Cantor model (Felsenstein, 2004) on an unrooted binary tree *T*, with *n* leaves. Each of the 2*n* – 3 edges of *T* have length λ, and thus the total tree length is *ξ* = λ(2*n* – 3). When *n* is large and *ξ* ⩽ 1, the expected number of substitutions per site is ⩽ 1; the number of edges on which a site changes state is approximately Poisson distributed with mean *ξ*; and the probability of more than one change on an edge is low, meaning multiple changes at a site occur on distinct edges. Though these conditions will generate alignments dominated by “perfect” sites exhibiting no homoplasy, a few sites may exhibit homoplasy even with *ξ* ⩽ 1, which motivates the term “near-perfect”. Under these conditions, tree reconstruction methods will tend to infer relatively unresolved trees unless the number of sites is very large.

Rare sites that exhibit homoplasy can introduce false positive splits on the inferred tree (Fig. 1). A naïve argument using Equation 1 might suggest that *FP_T_* would depend on *ξ* roughly as *O*(*ξ*^2^)/*O*(*ξ*) = *O*(*ξ*), namely the ratio of the expected numbers of sites having changes on two edges (i.e., those that are potentially homoplastic and misleading) to those sites having only a single change (those that are reliable), for sufficiently small *ξ*. But because only one-third of those two-edge sites are actually homoplastic in a JC69 model,

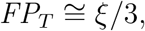

which implies *FP_T_* is small when *ξ* is small enough (e.g., *FP_T_* < 0.05 whenever *ξ* < 0.15).

This approximation can be improved further by recognizing that not all two-edge homoplastic sites induce false positives, depending on their position in the true tree (Fig. 1). Given the evolutionary model, the probability that *k* perfect sites, and another site *f* that has evolved with two edge changes will produce a “false positive” under MP is denoted 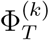 (Theorem 1 above). Because this probability is often less than one, *FP_T_* can remain below 0.05 at higher values of *ξ* than the naïve argument suggests.

If the true tree were known with some precision, the first part of Theorem 2 could be used directly to calculate false positive rates. However, in the “near-perfect” parameter space of large *n* and *ξ* ⩽ 1, estimates of the true tree are likely to be only partially resolved (Fig. 2). We therefore derive the expected false positive rate for a distribution, 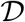, of randomly generated trees of size *n*, 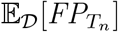, generated from parameters based on the inferred tree. In the remainder of this paper, the “expected false positive rate” will generally refer to 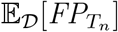. We assume that 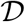 is usually either a “proportional-to-distinguishable-arrangement” (PDA) or Yule-Harding (YH) distribution (Aldous, 2001), but also consider the two extreme cases of completely unbalanced (caterpillar) trees, and completely balanced (symmetric) trees. Unlike PDA and YH trees, these last two have a constant tree shape (with random leaf labels). From the second part of Theorem 2, we see that, for a JC69 model and trees inferred with MP, the following approximation holds increasingly well as *n* increases:

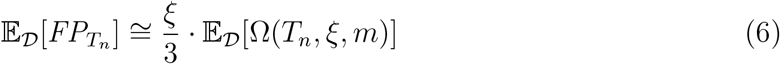

given the assumption that *ξ* is sufficiently small and the number of sites does not grow too quickly with the size of the tree. The function Ω(*T_n_*, *ξ*, *m*), defined in Materials and Methods, is monotonically decreasing in *ξ* and *m*, and depends on the shape of *T*. Simulations indicate that the approximation is close for *ξ* ⩽ 1 (Fig. 3), but if many equally parsimonious trees are present, the search algorithm should take a strict consensus of a broad sample of those solutions (Fig. S3). 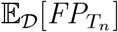 is better on average for PDA than YH trees, and both are bounded between a theoretical worst case error rate for symmetric and best case error rate for caterpillar trees. In fact, the expected false positive rate for the latter is just 4/(3*m*) in the limit of large *n*, which is independent of *ξ*.

### Robustness to Violation of Assumptions

Violations of assumptions tend to increase the expected false positive rate above the predictions of Equation 6. For example, adding edge length (EL) variation or across-site-rate (ASR) variation increases 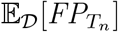 (Figs. 2, 4 and Fig. S4). The difference between predicted 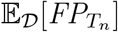 based on Eqn. (6), with no edge length variation, and simulation-based estimates with edge length variation included is small when *ξ* << 1 but increases substantially as *ξ* increases. When edge length variation is large (gamma shape parameter *α_e_* = 0.1), there is no longer a local maximum value of 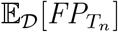 around *ξ* = 0.1; instead, 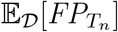 increases monotonically with *ξ* and eventually exceeds 5% for the simulated dataset sizes. The impact of ASR variation is deleterious at all values of *ξ*, but even when ASR variation is large (gamma shape parameter *α*_ASR_ = 0.1), the false positive rate remains slightly below 5% for simulated dataset sizes in the absence of EL variation (Fig. S4).

**Fig. 4.**
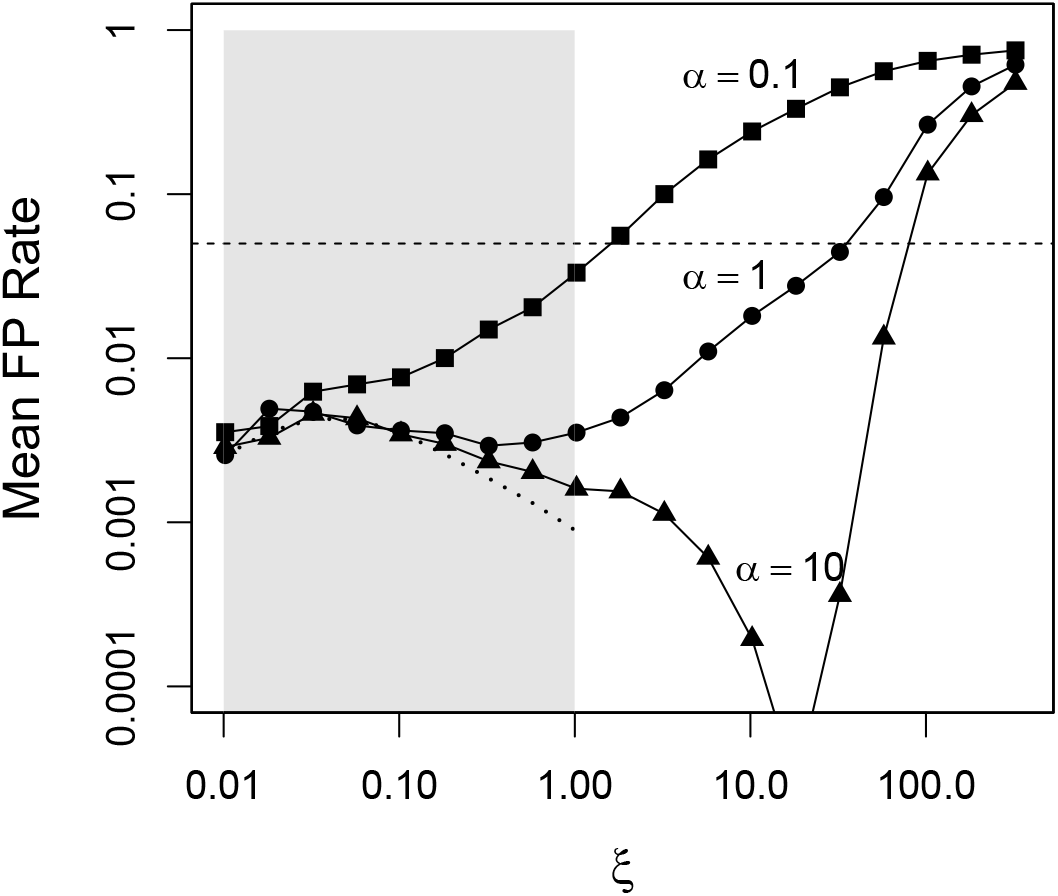
Effect of edge length variation on expected false positive rate for different values of the shape parameter of the edge length gamma distribution, *α_e_*. Smaller values of *α_e_* correspond to higher rate variation. ASR variation is assumed absent. The dashed curve is the prediction from Eqn. (6), in which both sources of variation are absent. Simulation conditions assumed PDA trees with *n* = 513, *m* = 1000, 1000 replicates, 100 subreplicates. Gray rectangle shows “near-perfect” values of *ξ* ⩽ 1.

Departure of the substitution model from the JC69 model assumed in the “near-perfect” zone can also increase the expected false positive rate. For example, a strong transition–transversion bias increases 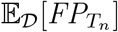 substantially, though it still remains well below 5% under our typical simulation conditions when *ξ* ⩽ 1 (Fig. S5).

Thus, the near-perfect tree length of *ξ* ⩽ 1 is a region in which rate variation appears to have less of an impact on false positive rates than when tree lengths are longer. This suggests that the definition of near-perfect zone in practice can include substantial rate variation as long as *ξ* ⩽ 1.

### Expected False Positive Rates in Virus Phylogenies

We estimated key parameters from the trees and underlying data for 11 empirical virus phylogenies (Table 1, S1) and used simulation to estimate expected false positive rates (Figs. 5, S6). The studies span a wide range of tree size and resolution and alignment length, and their tree lengths span three orders of magnitude. Seven of these viruses fell within the “near-perfect” tree length zone of *ξ* ⩽ 1.0, and five of those had 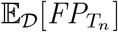 below 5% for two different models of ASR variation (see Methods) and both random tree models. 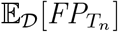 was uniformly lower for PDA vs. YH models, and for invariant sites vs. gamma models (Fig. S6). As expected, lower 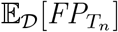 were generally observed for lower values of *ξ*.

**Fig. 5.**
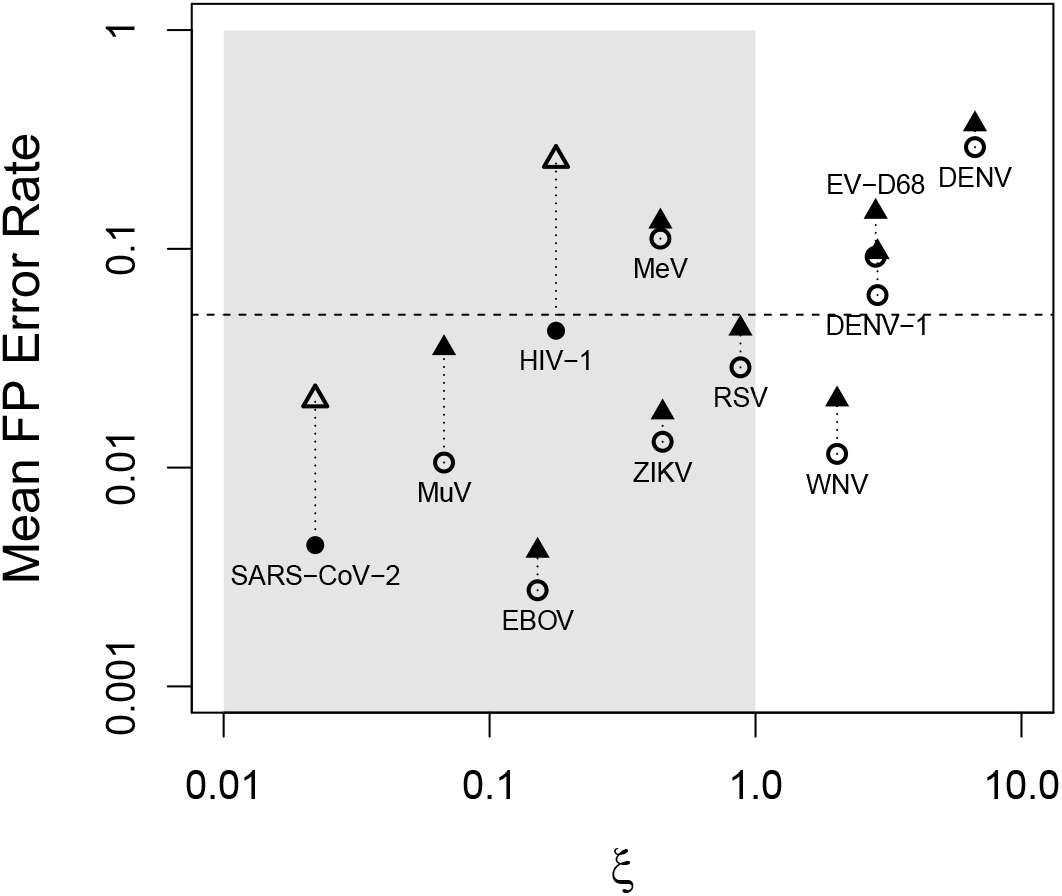
Expected false positive rates, 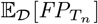, for 11 empirical virus phylogenetic datasets (Table 1) for maximum parsimony (MP) inference, estimated by simulation using parameters estimated from the data (Table S1). Abbreviations given in Table 1. Simulation experiments assumed ASR variation according to either an invariant sites model (circles) or a gamma distributed model (triangles) and a Yule–Harding random tree distribution (each point is mean of 500 replicates × 100 sub-replicates). Model point with higher likelihood is shaded. False positive rates assuming PDA random trees are uniformly slightly better (Fig. S6). The near-perfect zone of *ξ* ⩽ 1.0 is shaded. Horizontal dashed line indicates a 0.05 expected false positive rate.

Epidemics with young crown group ages on the order of years or decades (e.g., Zika virus, West Nile virus, and mumps virus) have an expected false positive rate below 5% expected in near-perfect trees, even though West Nile virus had a *ξ* slightly above 1. At the other extreme, dengue virus serotype 1, which does not represent a single epidemic, had a *ξ* > 1 and a correspondingly high expected false positive rate, and the phylogenetically more diverse dengue virus data representing all four serotypes had an even higher tree length and expected false positive rate.

The HIV-1 and measles virus trees both were outliers in having 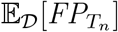 above 5%, even though their tree lengths were below one. These two trees had the fewest taxa (Table 1), possibly indicating sensitivity to the assumption of large *n* in our results. Moreover, the HIV-1 tree was constructed with the fewest sites (representing only a single partial gene), which affects accuracy through the Ω(*T_n_*, *ξ*, *m*) term in Eqn. 6. It may also lead to a poor estimate of the ASR gamma shape parameter, though 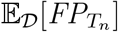 was about 5% when using the invariant sites model.

### Extension to Maximum Likelihood (ML) Inference

Theoretical results hint that ML and MP should reconstruct the same tree under

“near-perfect” assumptions. For example, ML provably converges to MP when there are enough constant characters in an alignment, a condition similar to *ξ* << 1 (Tuffley and Steel, 1997, Thm. 3). Further arguments presented in the SI support this conjecture.

We used simulation to check how well Equation 6, derived for MP, predicted the expected false positive rate under ML inference in the near-perfect zone. Simulations with *ξ* ⩽ 1, a JC69 model, and no edge length or ASR variation, with trees inferred by IQTree2 (Minh et al., 2020) under the same model, are close to the equation’s predictions (Fig. S7). Nonetheless, some differences were observed, which tended to imply better accuracy for MP. These differences could largely be attributed to technical or implementation issues. First, the computational expense of ML searches makes it tempting to undertake fewer replicate searches for local optima, but this was as critical to improve the fit to Equation 6 for ML as it was for MP (Fig. S7). Second, ML programs set hard numerical lower bounds strictly greater than zero on edge lengths, often (by default) on the same order as 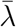, so these must be reset downward to obtain correct tree likelihoods (Morel et al., 2020). Finally, inferred edge lengths that are larger than these programs’ lower bounds but still smaller than about 1/*m* tend to be included in the ML tree despite weak evidence (IQTree2 issues a warning about this). We saw this in ML searches roughly when *ξ* ⩾ 0.1, when three-state sites become more common in alignments than they were at lower values of *ξ*. Even without homoplasy, ML tends to over-resolve trees in a way that elevates 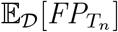. By collapsing short edge lengths inferred by ML to be less than 1/*m*, this behavior can be mitigated (Fig. S7).

In general, ML is expected to be more accurate than MP under more realistic model conditions and higher rates, something we observed commonly in simulations in which *ξ* > *ξ*\*. However, simulations also suggest that in the near-perfect zone, MP can achieve an 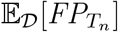 comparable with ML but with much faster running times.

### Accuracy and Support in Near-perfect Trees

False positive “accuracy”, defined as 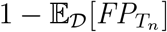, is very high in the near-perfect zone of small tree lengths, whereas conventional support values are quite variable in this zone under the same simulation conditions (Fig. 6). At very low *ξ*, the average MP bootstrap support is about the theoretically expected 64% for a single nonhomoplastic substitution supporting an edge (Felsenstein, 1985). Model-based support measures had higher values, with aBayes (Anisimova et al., 2011) being greater than ultrafast bootstrap (Hoang et al., 2018), which, in turn, was greater than SH-aLRT (Guindon et al., 2010), but only aBayes was close to 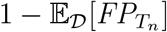 across the range of tree lengths in the near-perfect zone. Notably, aBayes is the only one of the four metrics that is not based on resampling.

**Fig. 6.**
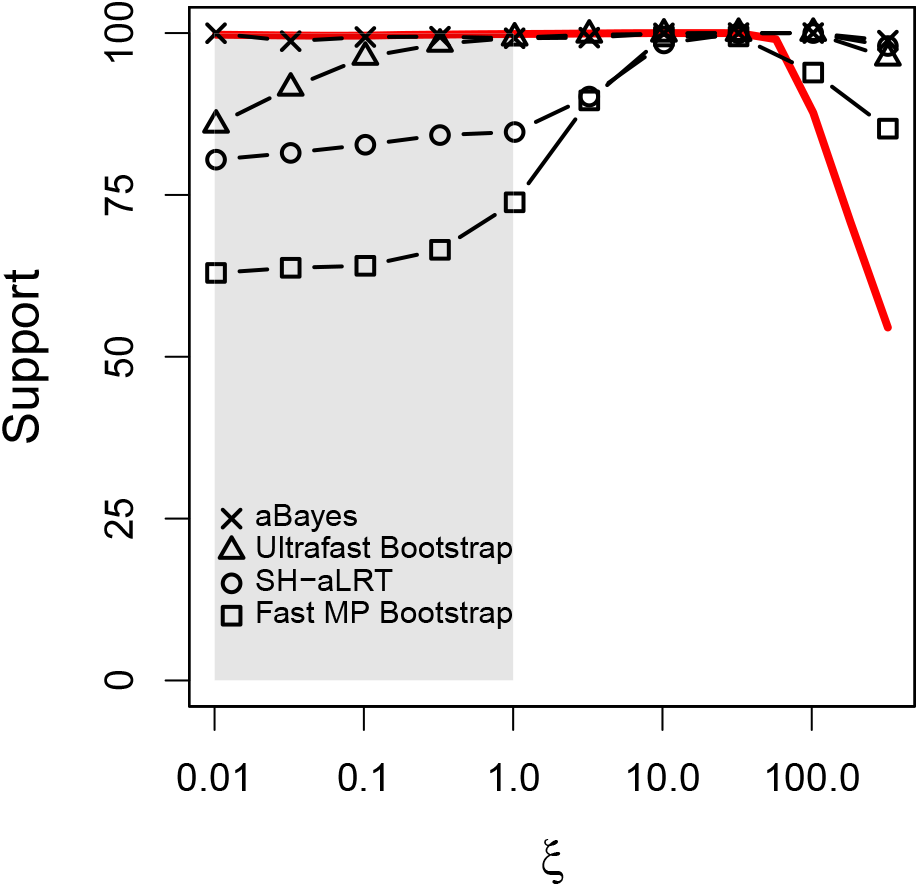
Statistical support measures compared to expected false positive accuracy. The red curve is the mean value of 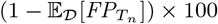 in simulations. The near-perfect parameter space is shaded.

We explored other factors impacting support in the boundary case of perfect trees. For sequence length, we computed standard support metrics in an ML framework in perfect four-taxon datasets, in which each branch was defined by a single change, and alignments range between 40 nt and 30,000 nt (Fig. S8). As observed for MP, non-parametric ML bootstrap support is approximately 63%, regardless of sequence length, in accordance with theoretical predictions (Felsenstein, 1973). Of the ML model-based support metrics, aBayes provided higher values than ultrafast bootstrap and SH-aLRT, both of which rely on bootstrap resampling. The aBayes support reached ⩾95% for alignments as short as 100 nt, which tracked the full Bayesian posterior support estimates that had support ⩾95% in alignments as short as 60 nt. The discrepancy between the Bayesian estimates and those that use bootstrap resampling, in light of our other results, suggests that resampling methods used in the presence of splits defined by only a single informative site may fail to integrate relevant information about low tree lengths.

On the other hand, in perfect trees from 8–128 taxa, in which the mean edge length remained the same (but therefore *ξ* grew with *n*), mean SH-aLRT and aBayes support was unchanged, but mean ultrafast bootstrap support increased (Fig. S9).

## Discussion

In this paper, we study a “near-perfect” parameter space for phylogenetic inference on large trees with small tree lengths and no rate variation within or between sites or edges. The “near-perfect” tree length of *ξ* ⩽ 1 means that few sites exhibit homoplasy and, for MP inference, the false positive rate can be much better than the false negative rate and well under 5% for typical datasets with thousands of sites. The near-perfect conditions defined here to allow mathematical derivations appear to be sufficient but not necessary. For example, with no rate variation, the false positive rate can be very good even when *ξ* > 1 (Fig. 2A, S5), and, if *ξ* < 1, a substantial level of rate variation can be present without elevating the false positive rate by nearly as much as when *ξ* > 1 (Fig. 2, 4, S4).

The second case is clearly more relevant in real-world data. The 11 empirical virus datasets all had substantial rate variation and showed a general increase in false positive rate with *ξ*, with almost all rates below 5% occurring when *ξ* ⩽ 1, much like the predicted patterns seen in Fig. 2B and Fig. 4. This accords with our simulation results suggesting that the good “near-perfect” false positive rates may emerge even when relaxing the strict near-perfect assumption of no rate variation—as long as *ξ* ⩽ 1.

These and many other empirical findings about RNA virus phylogenies sampled intensively in epidemics postdate much of the extensive body of other work on accuracy and support in phylogenetics. Not surprisingly, little note has been made about the stark contrast between false positive and false negative rates in phylogenies in which tree length is well below the optimal tree length for “overall accuracy”, since published examples have been relatively rare. The goal of much of the field of phylogenetics is, after all, to maximize tree resolution, even if this effort requires adding (or switching to) sequence data with more variation and thus longer tree lengths.

Because “near-perfect” datasets reflect a combination of the number of taxa and sites, evolutionary rate and time parameters, and assumptions about the substitution model, they also implicitly reflect sampling of the true tree, which is particularly relevant in epidemic trees in which sampling is far below disease incidence. Sampling can continue over time, increasing *n*, and the viruses continue to evolve over time, increasing the depth of the tree. Both of these increase *ξ* but in different ways; therefore, it is possible for the same RNA virus to have near-perfect and not near-perfect datasets depending on the study. For example, the SARS-Cov2 dataset we included had *n* = 583 and *ξ* = 0.02, well within the “near-perfect” zone, but a much more intensively sampled tree over a longer period of time (Lanfear, 2020) with *n* = 147156 has a tree length of *ξ* = 3.89 (after collapsing any edges with λ ⩽ 1.1 × 10^−6^), which is remarkably small for such a large tree but lies just outside our definition of near-perfect.

Other mathematical results on phylogenetic accuracy have largely focused on either the limiting case of infinite sequence length (“consistency”), or the number of sites needed for accurate inference (the “sequence length requirement”). For MP, for example, the shortest edge length is critical and 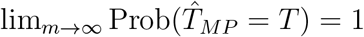 as long as λ_min_ > *ξ*^2^/(1 – *ξ*) (Steel, 2000, Thm. 1(A)). More generally, let *m*′ be the number of sites needed for 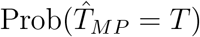 to exceed some fixed required accuracy. For the neighbor-joining method *m*′ grows exponentially with *n* (Lacey and Chang, 2006); for ML, *m*′ is polynomial or better in *n*, depending on edge lengths (Roch and Sly, 2017). Moreover, *m*′ also grows as 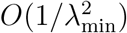 for ML and some more ad hoc estimators (Erdos et al., 1999; Roch, 2019), implying again that short edges tend to degrade accuracy when accuracy is defined in terms of total agreement between *T* and 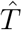, unlike here.

A cryptic factor affecting the false positive rate is tree shape. Highly asymmetric trees have better expected false positive rates than highly symmetric trees, because expected path lengths are longer and it is harder to induce false positive splits by chance (Fig. 1). Thus, a random sample of PDA trees will have a better 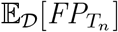 than more symmetrical YH trees. Differences in tree shape among RNA virus phylogenies have long been noted (Grenfell et al., 2004), such as the typically more asymmetric influenza trees.

Perfect and near-perfect phylogenies have been studied as discrete optimization problems (Gusfield, 1997; Fernandez-Baca and Lagergren, 2003) in which the goal is to find an optimal tree when, at most, some small number of sites exhibit homoplasy. Little of this work has considered accuracy per se, but Gronau et al. (Gronau et al., 2012) highlighted the connection between short edge lengths and false positives, and developed a “fast converging” algorithm (i.e., having an *O*(poly(*n*)) sequence length requirement) that returns a tree with short edges collapsed when they do not meet a threshold probability of being correct, thus minimizing false positives. The connection between this tree and those built by more conventional methods is unclear, but it may be a promising approach for trees in the near-perfect zone.

Model-based phylogenetic inference methods such as ML and Bayesian inference are generally regarded as theoretically superior to MP, especially for datasets that fit substitution models much more complex than our “near-perfect” JC69 model with no rate variation. Though our mathematical results for expected false positive rates were derived for MP, there is both relevant theory and considerable simulation evidence to suggest that in the near-perfect zone, the ML expected false positive rate is approximated by the MP theory, both in terms of its absolute value and its shape as a function of tree length. As *ξ* increases, especially above *ξ*\*, ML consistently has better accuracy than MP, but we conjecture that the false positive rates of MP and ML differ much less *ξ* as gets very small. Further work is needed to test this conjecture.

The connection between the false positive rate as a measure of accuracy and conventional measures of phylogenetic support appears to be sensitive to the choice of support method when *ξ* << 1 (Fig. 6). The aBayes method corresponds well to what is implied by 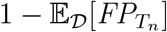, but resampling methods using either likelihood or parsimony correspond less well. The connection between phylogenetic accuracy and support in frequentist and Bayesian settings has been studied in detail (Felsenstein, 1985; Hillis and Bull, 1993; Felsenstein and Kishino, 1993; Efron et al., 1996; Susko, 2008, 2009; Alfaro and Holder, 2006; Simmons and Norton, 2014), but remains somewhat fraught. We hesitate to draw firm conclusions without a formal analysis of support in the “near-perfect” parameter space but note the variability in support estimates we found (Fig. 6).

The low false positive rate in near-perfect trees suggests that phylogenies describing viral epidemics in this zone can be interpreted directly without defaulting to identifying clades with strong support values. Frequent convergent evolution, and recombination in positive-strand RNA viruses, can complicate phylogenetic inference and may increase the false positive rate in real-world trees (Morel et al., 2020). Nonetheless, if individual clade support needs to be invoked, we recommend Bayesian approaches that do not rely on bootstrap resampling of sparse substitutions.

The benefit of real-time viral genomic sequencing for public health action became apparent during the 2014–2015 West African Ebola epidemic (Gire et al., 2014), and is a critical component of tracking the COVID-19 pandemic (Oude Munnink et al., 2020; Grubaugh et al., 2021). Consequently, the viruses responsible for these diseases, Ebolavirus and SARS-CoV-2, epitomize near-perfect phylogenetic trees in our analysis. We can expect a greater intensity of genomic sequencing accompanying future viral outbreaks, increasing the importance and relevance of near-perfect phylogenies.

## Acknowledgements

We gratefully acknowledge the authors from the originating laboratories and the submitting laboratories, who generated and shared via GISAID the SARS-CoV-2 genomic sequence data on which this research is based. A complete list acknowledging the authors who submitted data analyzed in this study can be found in Data S1. MJS thanks the University of Arizona’s HPC facility, Bio5 Institute, and Rod Wing’s lab for computing support. JOW was supported by an NIH-NIAID R01 AI135992.

## Supplementary Information

### Mathematical Results and Proofs

#### Definitions and preliminary observations

Let *B*(*n*) denote the set of unrooted binary phylogenetic trees with leaf set [*n*] = {1, 2, …, *n*}. Thus, *B*(*n*) = (2*n* – 5)!!. Note that a tree *T* ∈ *B*(*n*) has 2*n* – 3 edges. Consider a Jukes-Cantor model in which all edges have the same value of λ. Thus, the probability of a state change between the endpoints of an edge *e*, denoted *p_e_*, is given by *p_e_* = *p* where 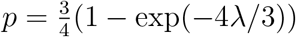.

A *character* refers to the assignment of states to the taxa at a given site of an alignment. Let us say that a character evolves ‘perfectly’ on *T* if there is a single change of state across one interior edge (say *e*) and no change of state any other edge of *T*. The probability that this occurs for any given interior edge *e* is: 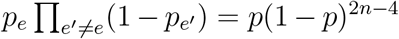. Note that this probability does not depend on the particular choice of *e* or on the shape of *T*. Note also that a perfectly evolved character has at least two species present in each state (otherwise the change would have been on a pendant edge of *T*).

We will say that a character *f* evolves on *T* with *c edge changes on e*_1_, …, *e_c_* if state changes occur on edges *e*_1_, …, *e_c_* and on no other edge of *T*. More briefly, we say that *f* evolves on *T* with *c* edge changes if *f* evolves with *c* edge changes for some set of *c* distinct edges of *T* (mostly we will deal with the case *c* =2). The probability that a character evolves on *T* with 2 edge changes on *e*_1_, *e*_2_ is *p*^2^(1 – *p*)^2*n*–5^, while the probability that a character evolves on *T* with 2 state changes is 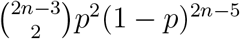.

We pause here to make an observation: If a data set consists of characters each of which have evolved on *T* with either 1 or 2 edge changes, then the maximum parsimony tree(s) and the maximum compatibility tree(s) for this data set will be exactly the same. Moreover, if there are sufficiently many constant characters, then any maximum likelihood tree will also be one of these trees.

Recall that a *split* refers to a bipartition of the leaf set [*n*] into two non-empty subsets (and splits are induced by binary characters). A character that has evolved perfectly on *T* produces a split, and these splits (across a set of perfectly evolved characters) are compatible and so form a (generally unresolved/non-binary) tree on leaf set [*n*].

Now suppose that *m* characters evolve on *T* and suppose that, of these *m* characters, *k* of them are perfectly evolved on *T* (note that more than one of these characters may correspond to the same split of *T*).

Next, consider a single additional character *f* which has evolved on *T* with two edge changes on *e*_1_, *e*_2_ (there is no restriction that these are interior edges). The probability that this character *f* is a binary character is 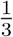 (as noted earlier). Moreover, in that case, the bipartition induced by *f* corresponds to a split of *T* if and only if *e*_1_ and *e*_2_ are adjacent.

The question now arises as to whether we can discover that *f* gives a false split of *T* (without knowing *T*), just on the basis of the other *k* perfectly-evolved characters.

We next introduce some further notation. We will let *ξ* denote the expected number of state changes in the tree *T* per character, under the model described. Thus *ξ* equals the per-edge rate λ times the number of edges of *T*, and so

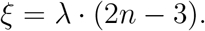

We will also use the following standard notation throughout: we write *f*(*n*) ~ *g*(*n*) if the ratio *f*(*n*)/*g*(*n*) tends to 1 as *n* becomes large.

Given a tree *T* ∈ *B*(*n*), let *d_T_*(*e*_1_, *e*_2_) denote the number of edges of *T* that lie strictly within the path between *e*_1_ and *e*_2_ (i.e. excluding *e*_1_ and *e*_2_). Thus, *e*_1_ and *e*_2_ are adjacent if and only if *d_T_*(*e*_1_, *e*_2_) = 0

Next, given *T* ∈ *B*(*n*), let *φ_T_* = (*φ_T_*(0), *φ_T_*(1), …, *φ_T_*(*n* – 3)) where *φ_T_*(*i*) is the number of (unordered) pairs of edges {*e*, *e*′} of *T* for which *d_T_*(*e*, *e*′) = *i*. Thus *φ_T_*(0) = 3(*n* – 2), however the other values comprising *φ_T_* depend on the shape of *T*. In particular, for a complete balanced binary tree with *n* = 2^*h*^ leaves we have *φ_T_*(*i*) = 0 for all *i* ⩾ 2log_2_(*n*) – 2, while for a caterpillar tree with the same number of leaves we have *φ_T_*(*i*) > 0 for all *i* ⩽ *n* – 3.

Note that the sequence *φ_T_* is a topological invariant (i.e. it depends only on the unlabelled shape of the tree) and does not depend on any other parameters mentioned above. Clearly, for any *T* ∈ *B*(*n*) we have:

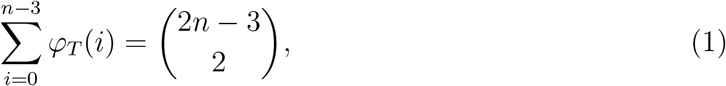

since both sides of this equation count the number of pairs of edges of *T*. For *i* between 1 and *n* – 3, we will let

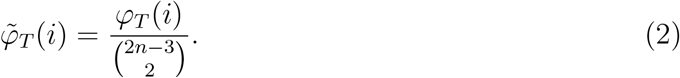

Thus, Eqn. (1) translates to the identity 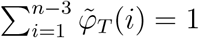. Thus 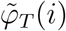 is the probability that a pair on edges (selected uniformly at random from all pairs) has exactly *i* edges lying on the path strictly between the two selected edges.

Next we state a simple combinatorial result that will be useful in the proof of the first theorem.

##### Lemma 1

Let *f* be a binary character that has evolved on *T* by 2 edge changes on *e*_1_ and *e*_2_, and let *f*′ be a character that has perfectly evolved on *T* by a change on a single interior edge *e*. Then *f* and *f*′ are compatible (i.e. induce compatible splits) if and only if *e* does not lie on the path in *T* that is strictly between *e*_1_ and *e*_2_ (i.e. the red edges in Fig. 1).

*Proof*: First suppose that *e* lies on one of the red edges shown in Fig. 1, say between *t_i_* and *t*_*i*+1_ for *i* ∈ {1, · …, *r* – 1}. Let *x_j_* be a leaf of *t_j_* for *j* ∈ {0, *i*, *i* + 1, *r* + 1}. Then if *f* is a binary character arising from changes on (just) *e*_1_ and *e*_2_ then *f* splits the set {*x*_0_, *x_i_*, *x*_*i*+1_, *x*_*r*+1_} as *x*_0_*x*_*r*+1_|*x_i_x*_*i*+1_ while *f*′ splits this same set as *x*_0_*x_i_*|*x*_*i*+1_*x*_*r*+1_. Since these two partial splits are incompatible, so too are *f* and *f*′.

Next suppose that *e* lies in one of the green subtrees of Fig. 1 (including the stem edge of that subtree), say subtree *t_i_* for *i* ∈ {0,1, …, *r* + 1}. We consider the possible cases: (1) *e* ∈ {*e*_1_, *e*_2_}, or (2) *e* is an edge of *t*_0_ or *t*_*r*+1_ or (3) *e* is an edge of *t_i_* or its stem edge for *i* ∈ {1, …, *r*}. Let *X_i_* ⊂ [*n*] denote the subset of leaves of *T* that are leaves of *t_i_*.

In case (1), it suffices by symmetry to consider the case *e* = *e*_1_. Then *f*′ induces the full split *X*_0_|([*n*] – *X*_0_) while *f* induces the full split (*X*_0_ ∪ *X*_*r*+1_)|([*n*] – (*X*_0_ ∪ *X*_*r*+1_)) and these two splits are compatible, since the set on the left of the first split is contained in the set on the left of the second split.

In case (2), it suffices by symmetry to consider the case where *e* is an edge of *t*_0_. In that case *f*′ induces a full split of the for *Y*|([*n*] – *Y*) where *Y* ⊆ *X*_0_, and this split is again compatible with the full split (*X*_0_ ∪ *X*_*r*+1_)|([*n*] – (*X*_0_ ∪ *X*_*r*+1_)) induced by *f* since *Y* is a subset of the left part of this split.

In case (3), suppose that *e* is an edge of *t_i_* or its stem edge. In that case, *f*′ induces the split *W*|([*n*] – *W*) where *W* ⊆ *X_i_*, and since *X*_0_ ∪ *X*_*r*+1_ ⊆ [*n*] – *W* it follows that *f* is compatible with *f*′. This completes the proof of Lemma 1.

Next, let 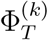 be the probability that a character *f* that has evolved on *T* with 2 edge changes has the following three additional properties:

The point of these conditions is that we might add the split corresponding to *f* to the other *k* compatible spits which still be compatible, but create a ‘false positive’ split in the resulting tree (note that we do not know a-priori which splits are the perfectly evolved ones, as we are assuming that *T* is not known).

We now present an exact expression for 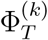. Firstly, let 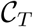 be the collection of all (unordered) pairs of non-adjacent edges in *T*. Thus 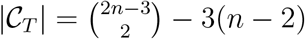. For 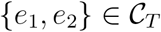, let

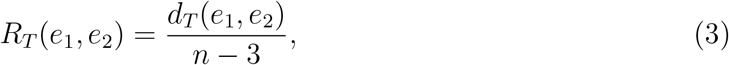

Thus *R_T_*(*e*_1_, *e*_2_) is the proportion of interior edges of *T* that lie between two non-adjacent edges *e*_1_ and *e*_2_.

**Fig. 1.**
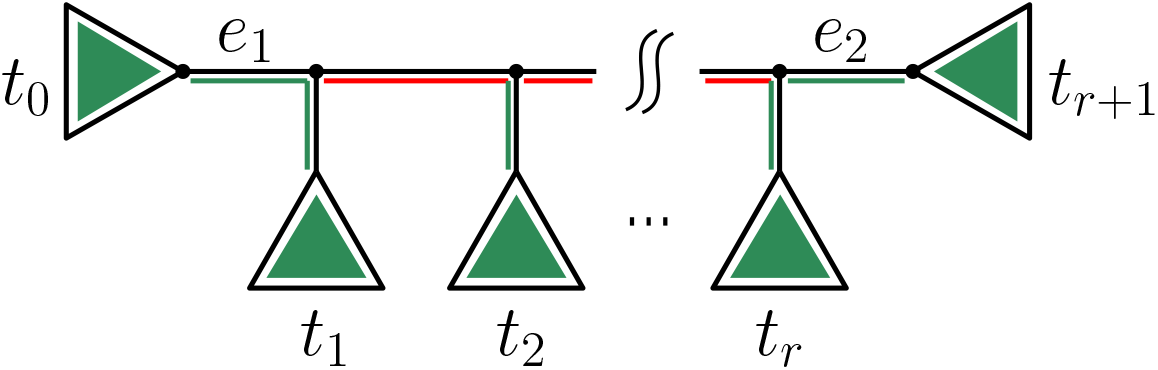
A representation of *T* determined by the pair {*e*_1_, *e*_2_} on which changes of state occur for character *f*. When *f* is a binary character, a perfectly evolved character *f*′ will be compatible with *f* provided *f*′ corresponds to a change of state on an edge in the green portions of the tree; otherwise, if the change is on one of the *r* – 1 edges marked in red, the two characters will be incompatible.

Next, let *μ_T_* denote the average value of *d_T_*(*e*, *e*′) over all pairs of edges {*e, e*′} sampled from *T*. That is:

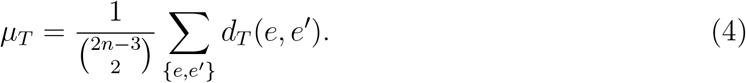

##### Theorem 2

For each *T* ∈ *B*(*n*), and *k* ⩾ 1 we have:

i. 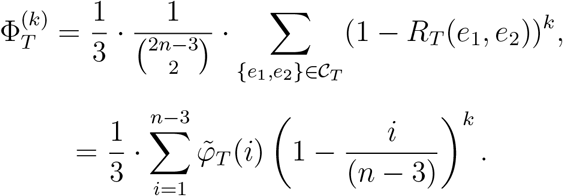
ii. 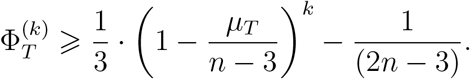

*Proof: Part (i):* Under the Jukes-Cantor model, the (conditional) probability that a randomly evolved character *f* is binary, given that is has evolved on *T* with 2 edge changes is 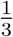. To see this, let *e*_1_, *e*_2_ denote the two edges on which there are state changes. Then if the state at the left-hand end vertex of *e*_1_ (as shown in Fig. 1) is *α* and the state on the right end vertex of *e*_1_ is *β* then for *f* to be a binary character we require *e*_2_ to involve the transition from *β* back to *α* (which is one of the three possible other states that *β* can change to on *e*_2_, and each change has equal probability under the Jukes-Cantor model).

Now consider a single random character *f*′ that has perfectly evolved on *T*. Since (a) each of the *n* – 3 interior edges of *T* has the same probability of being the ‘change edge’ associated with *f*′, and (b) *f*′ is compatible with *f* if and only if the associated change edge for that character does not lie on the path that is (strictly) between *e*_1_ and *e*_2_ (by Lemma 1) and (c) the proportion of interior edges of *T* that do not lie strictly between the edges *e*_1_ and *e*_2_ is 1 – *R_T_*(*e*_1_, *e*_2_), it follows that the probability that *f*′ is compatible with *f* is 1 – *R_T_*(*e*_1_, *e*_2_).

Thus, for *k* characters that have independently perfectly evolved on *T* the probability that each of them is compatible with *f* is (1 – *R_T_*(*e*_1_, *e*_2_))^*k*^.

Finally, each choice for *f* across all pairs of edges of *T* (nor just the non-adjacent pairs) has equal probability, and if we let *η*(*e*_1_, *e*_2_) = 1 if {*e*_1_, *e*_2_} are not adjacent, and 0 otherwise, then the probability that *f* satisfies properties (C-i), (C-ii) and (C-iii) above is the average value of 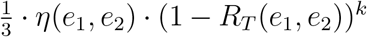 across all 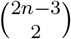 pairs of edges {*e*_1_, *e*_2_} of *T*, which gives the first expression in Theorem 2(i), and the second expression follows directly.

*Part (ii*). Let 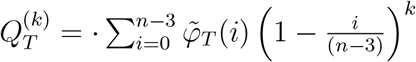. Then 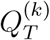 is the expected value of (1 – *R_T_*(*e*_1_, *e*_2_))^*k*^ when a pair of edges of *T* is selected uniformly at random from the set of all such pairs. Since *f*(*x*) = (1 – *x*)^*k*^ is a convex function, Jensen’s inequality then gives:

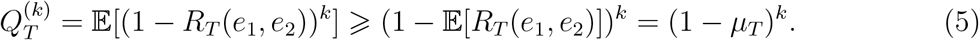

Now 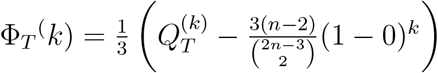, since there are 3(*n* – 2) pairs of adjacent edges in *T*, and 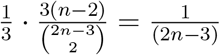, and so applying this to the Eqn. 5 gives the result claimed.

##### Remarks

Theorem 2(i) shows that for fixed *k* and *n*, the shape of *T* plays a significant role in determining 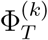; in particular, pectinate trees (such as caterpillars) will have a smaller value of 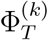 than more balanced trees. Indeed, it is possible to exactly calculate the value of 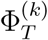 for the two extreme families: caterpillars and fully-balanced trees to determine the extent of this dependence, as we describe in the next section. Notice also that the exact computation of 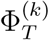 for a given tree *T* with *n* leaves involves a summation of just *O*(*n*^2^) terms, so could be calculated fairly easily.

Notice also that in the special case where *k* =1, Theorem 2(i) simplifies (via Eqn. (1)) as follows.

##### Corollary 1

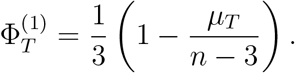

#### Exact values of 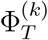 for classes of trees

##### Proposition 1

i. Let *T_n_* denote a caterpillar tree with *n* leaves. Then Φ_*T*_4__ = 0 and for *n* ⩾ 5:

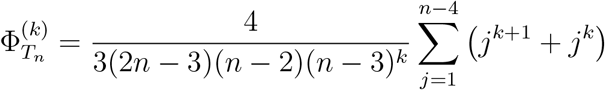 Thus, for fixed *k* we have:

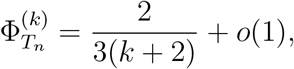

where *o*(1) is a term that tends to 0 as *n* grows.
ii. Let *T^h^* denote any tree in *B*(2^*h*^ + 1) obtained by attaching a leaf to the root of a complete balanced binary tree of height *h*. Then

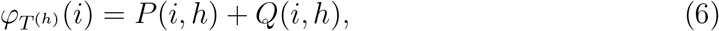

where:

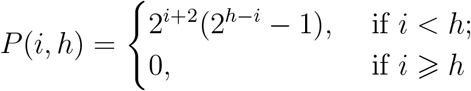

and

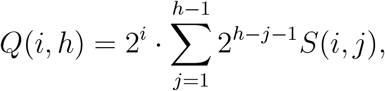

where

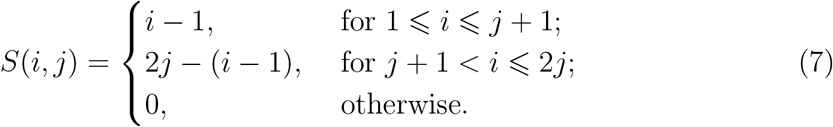
iii. Let 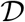 be a probability distribution on *B*(*n*). Then

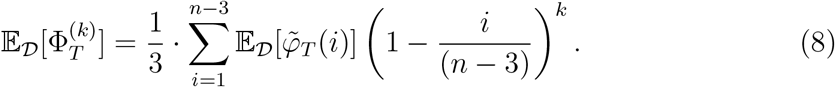 In particular, when 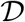 is the PDA distribution on *B*(*n*)

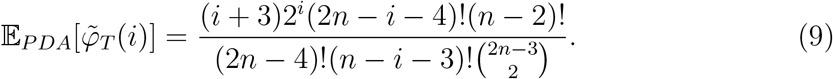

*Proof: Part (i):* For *n* ⩾ 4 any path of length ℓ in *T_n_* that lies strictly between two edges {*e, e*′} is a path of interior edges, and there are precisely four such pairs of edges that correspond to the same path. Moreover, the number of interior edge paths of *T_n_* of length *n* – 3 – *j* is *j* + 1 for each *j* between 0 and *n* – 4. Thus,

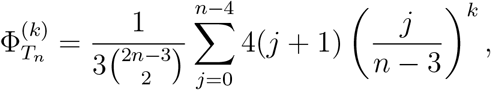

and straightforward algebraic manipulation leads to the stated expression. The last part of (i) follows from the asymptotic identity: 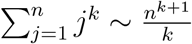.

*Part (ii*)

For *h* ⩾ 1, let 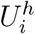 be the set of edges *e* of *T^h^*, excluding the stem edge, for which there are exactly *i* ⩾ 0 edges strictly between *e* and the stem edge. If we also let 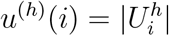 then it is easily seen that:

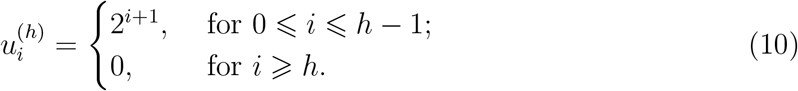

For *h* ⩾ 1 and *i* ⩾ 1, let 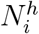 denote collection of pairs of edges in *T^h^* that are separated by exactly *i* edges, and let 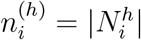. Thus, 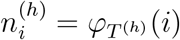. Observe that:

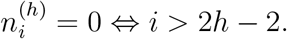

Notice also that deleting the stem edge of *T*^*h*+1^ (and its incident vertices) produces two copies of *T^h^* - we will call these *L* and *R* (‘left’ and ‘right’).

The set 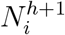 can be partitioned into three classes:

**Class 1:** A pair of edges consisting of an edge of *T*^*h*+x^ at distance *i* from the stem edge, together with the stem edge. This set has size 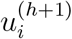, where 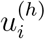 is as given above.
**Class 2:** A pair of edges that lie entirely within *L* or entirely within *R*. This set has size 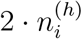.
**Class 3:** A pair of edges {*e*_1_, *e*_2_} with *e*_1_ in *L* and *e*_2_ in *R* with the distance between *e*_1_ and *e*_2_ being *i*.

A little care is required to enumerate Class 3. One subcase is that *e*_1_ is not the stem edge of *L* and *e*_2_ is not the stem edge of *R*. In that case, the number of choices is

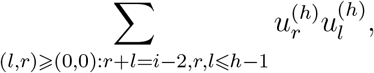

which equals the expression *Q*(*i, h*) given in Part (ii). The complementary subcase where *e*_1_ or *e*_2_ is the stem edge of *L* or *R* contributes

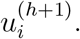

Combining the above gives the recursion:

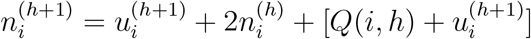

which simplifies to:

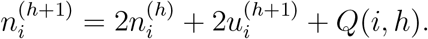

or equivalently,

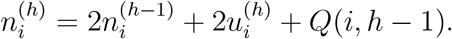

Solving this recursion (noting that 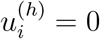 for *i* ⩾ *h*) leads to the expression for *ψ_T_*^(*h*)^ given in Part (ii).

*Part (iii):* Eqn. (8) follows by linearity of expectation (and interchanging the order of expectation operators).

The expression for 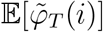 involves an argument in enumerative combinatorics. Let *N*(*n, k*) denote the number of (unordered) forests consisting of *k* rooted binary trees whose leaf sets are disjoint and contain a total of *n* leaves (we allow a single labeled leaf to be a rooted tree of size 1, otherwise the root of each tree has degree 2). It is easily seen that *N*(*n*, 2) is precisely the number of rooted binary trees (2*n* – 3)!!, since deleting the root of such a tree, produces a forest of two trees with disjoint leaf sets. It turns out there is an exact formula for *N*(*n, r*) (from ~1990):

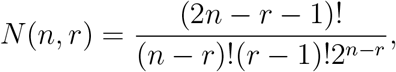

for *r* = 1, …, *n* (see e.g. (3), p. 105).

An *ordered forest* is a forest with a linear ordering on its component trees. If *O*(*n, r*) denotes the number of ordered forests consisting of *r* trees then

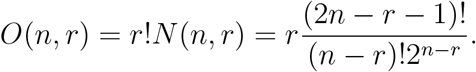

Notice that:

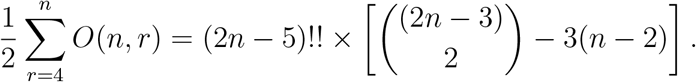

To see this observe that twice the right-hand side counts the number of pairs (*T*, (*e*_1_, *e*_2_)) where *T* ∈ *B*(*n*) and (*e*_1_, *e*_2_) is an ordered pair of non-adjacent edges. Notice that if we delete the path connecting *e*_1_ and *e*_2_ we obtain an ordered forest of rooted trees on the same leaf set as *T* (*cf*. Fig. 1). This provides a bijection between these two sets in which the number of strictly edges between *e*_1_ and *e*_2_ in *T* is equal to *i* – 3 where *i* is the number of forests.

Thus,

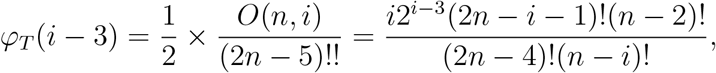

and rearranging, gives the result.

To illustrate Part (i), with *n* = 5 we have 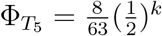. Notice that this converges to zero exponentially fast with *k* (and in general for fixed *n* this will be the case).

Also, observe that for general values of *n* and with *k* =1 (say) we have:

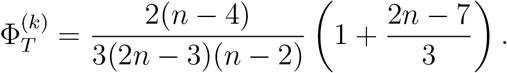

Thus, for the simulation involving trees with *n* = 500 leaves, if these were caterpillars (instead of YH trees), for *k* =1 we would expect 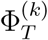 to be very close to 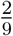 (in agreement with the asymptotic claim in the Part (i) of Proposition 1) which is lower than the simulated values on YH trees, as expected.

To illustrate Part (ii), *φ*_*T*^3^_(2) = *P*(2, 3) + *Q*(2, 3) = 2^4^(2^1^ – 1) + 2^2^(2 + 1) = 28. Notice also that for general *h* we obtain (as expected)

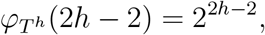

and *φ_T^h^_*(*i*) = 0 for *i* > 2*h* – 2.

Notice that for *h* large, *P*(*i, h*) is negligible relative to *Q*(*i, h*).

#### A lower bound on the expected value of 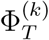 for the PDA and YH distributions

##### Proposition 2

Let *T* be a tree in *B*(*n*) sampled from the PDA or YH distribution. Then, for all *n* ⩾ 4 we have:

i. 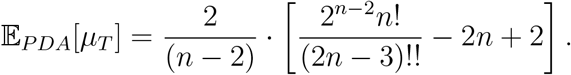 Moreover,

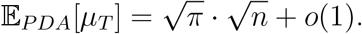
ii. 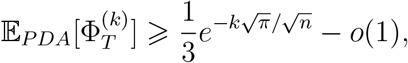

where *o*(1) is a term that tends to 0 as *n* grows.
iii. 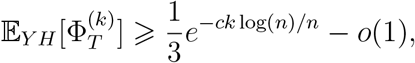

where *o*(1) is a term that tends to 0 as *n* grows, and *c* is a constant (independent of *n*).

*Proof: Part (i):* For *T* ∈ *B*(*n*) let *NA*(*T*) be the collection of (unordered) pairs {*e*_1_, *e*_2_} of non-adjacent edges of *T*.

The first step is to apply a classic technique in enumerative combinatorics; namely, we count a certain set Ω in two different ways, to obtain an equation. Here, we take the set Ω to be the collection of triples (*T*, *σ*, {*e*_1_, *e*_2_}) where *T* ∈ *B*(*n*), *σ* is a split of [*n*] that corresponds to an interior edge of *T*, and {*e*_1_, *e*_2_} ∈ *NA*(*T*) with the edge of *T* corresponding to split *σ* being strictly within the path connecting *e*_1_ and *e*_2_.

Summing first over all choices of *T* we have:

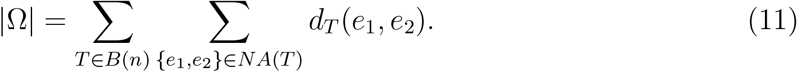

and so,

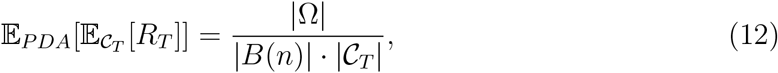

where 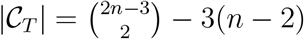 is the number of pairs of non-adjacent edges in *T*.

On the other hand, we can count | Ω| by first summing over all choices of the split *σ* (stratified by the size *k* of the smaller half of the split) and then counting for each such *σ* the number of *T* and {*e*_1_, *e*_2_}. This gives:

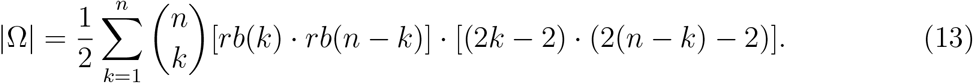

where *rb*(*m*) = (2*m* – 3)!! is the number of rooted trees on a leaf set of size *m*.

To see this, note that 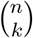 is the number of ways to partition [*n*] into a split *σ* of two sets of size *k* and *n* – *k*, and 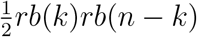 is the number of trees in *B*(*n*) that contain this split (the factor 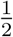 is to remedy the double counting that occurs). Finally, given *T* and *σ* there are now (2*k* – 2)(2(*n* – *k*) – 2) choices of {*e*_1_, *e*_2_} where *e*_1_ is an edge of the rooted subtree of *T* on the leaf set of size *k* and *e*_2_ is an edge of the rooted subtree of *T* of size *n* – *k*.

We will apply generating function techniques to calculate an exact expression for term on the right hand side of Eqn. (13). Notice can rewrite Eqn. (13) as follows:

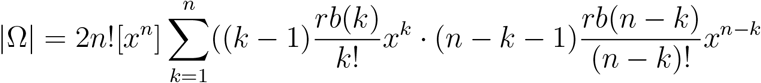

where [*x^n^*]*f*(*x*) denote the coefficient of *x^n^* in *f*(*x*), and hence, even more compactly,

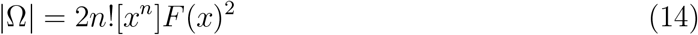

where

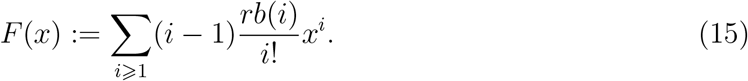

Let *rb*(*n*) = (2*n* – 3)!! (the number of rooted binary trees on a leaf set of size *n*) and let 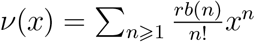 denote the associated exponential generating function. It is well known (see e.g. (3)) that

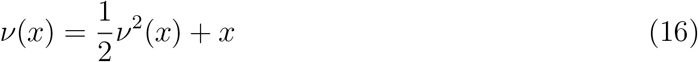

which has the unique solution

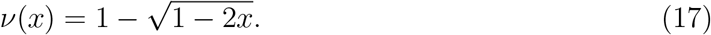

We will use the following relationships which follow easily from Eqn. (16):

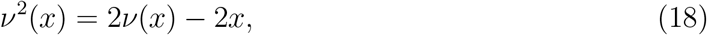

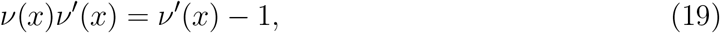

where 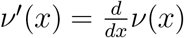.

Recall, *F*(*x*) from Eqn. (15). We have:

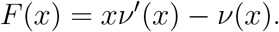

Thus,

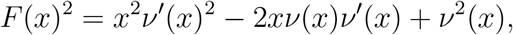

and this simplifies by Eqns. (18) and (19)) to:

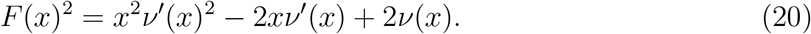

In order to determine the expression in Eqn. (14) we consider the coefficient of *x^n^* in *F*(*x*)^2^ (i.e. [*x^n^*]*F*(*x*)^2^) by considering the three terms on the right of Eqn. (20). For the first term of Eqn. (20), Eqn. (17)) gives: 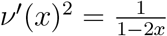 and so

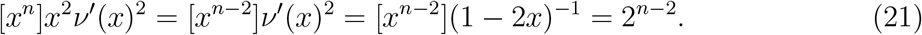

For the second term of Eqn. (20), we have:

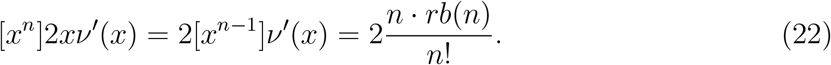

For the third term of Eqn. (20), we have:

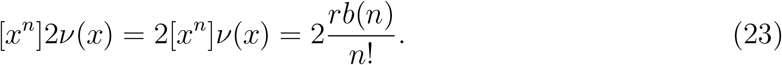

Substituting the expressions on the right hand side of Eqns. (18), (19) and (20) into the corresponding terms in Eqn (20) allows us now to determine the expression for |Ω| given by Eqn. (14), as follows:

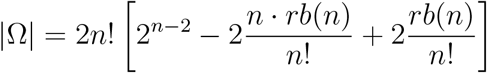

which simplifies slightly to:

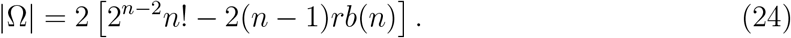

By substituting the expression for |Ω| given by Eqn. (24) into Eqn. (12) and rearranging terms, gives the claimed expression for 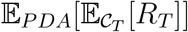.

The second claim in Part (i) of Theorem 2 follows from the asymptotic equivalence:

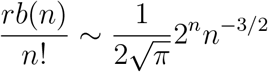

together with some standard algebraic manipulation.

*Part (ii):* We apply Theorem 2(Part (ii)) to Part (i) of the current theorem. WIth 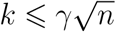 we obtain:

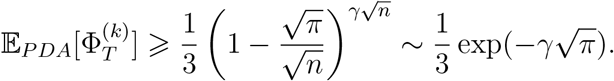

*Part (iii):* We apply Proposition 4 of (1) (where *β* = 0 corresponds to the YH distribution). This result shows that in a tree *T* ∈ *B*(*n*) sampled according to the YH distribution the maximum distance *D* from any leaf to the root is less than (4.31 + *ϵ*) log(*n*) with a probability converging to 1 as *n* grows. Now *μ_T_* is always less than twice the distance from any leaf in *T* to any other leaf in *T*, and this inter-leaf distances is bounded by 2*D*. The result now follows from Theorem 2(Part (ii)).

##### Remarks

Note that Parts (ii) and (iii) imply that for each fixed *k*, 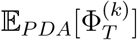 and 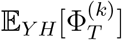 both converge to 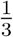 as *n* grows, in contrast to the caterpillar case of Part (i) of Proposition 1, where the limit is smaller (e.g. for *k* =1, the limit is 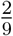).

Notice also that the lower bounds on 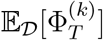 allow *k* to grow faster with *n* for YH trees than PDA (essentially because in a Yule tree, there are on average fewer edges on paths between leaves, so fewer ‘red’ edges on the path between *e*_1_, *e*_2_ (see Fig. 1). In this lower bound we are undercounting by ignoring the red portions to the left of *e*_1_ and right of *e*_2_ by considering only cases where *e*_1_ and *e*_2_ are both pendant edges (this undercounting is valid since we are stating a lower bound on 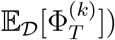.

#### The expected number of such false splits

Note that Theorem 2 describes the probability that a single character satisfies the three conditions (C-i)–(C-iii) above, conditional on this character having evolved on *T* with 2 edge changes (thus 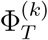 should be viewed as a conditional probability). This raises two further natural question: for a given character, what is the probability *p_T_* that such a character will evolve so as to satisfy conditions (Ci)–(Ciii)? And how many such characters should we expect to see? For these questions, the number *k* of perfectly evolved characters should now be treated as a random variable, *K*. Thus, let *K* be the random variable corresponding to the number of distinct (and nontrivial) splits generated from those characters (from within the *m* characters in total) that have perfectly evolved on *T*. The variable *K* lies between 0 and *m*. Let *μ_K_* be the expected value of *K*.

Conditional on *K* = *k*, the value *p_T_* is simply given (ignoring terms involving λ of higher than quadratic power) by:

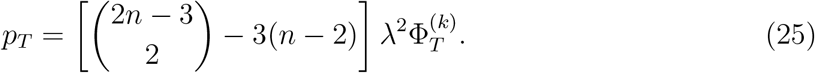

In particular,

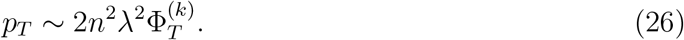

We have

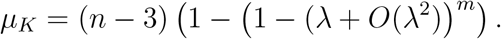

Provided that λ*_m_* << 1 we may approximate this last equation by:

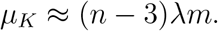

Thus, provided that *m* large and λ_*m*_ < 1 and using using *μ_K_* to estimate *K* (which is reasonable since *m* is large, and we are also assuming above that λ*n* << 1) we obtain:

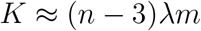

and solving for λ in this equation, and substituting the estimate of λ into Eqn. (26) gives:

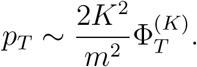

Let us now multiply through by *m* (the number of characters) to get.

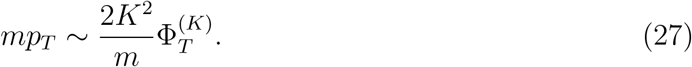

The term on the left (*mp_T_*) has a natural interpretation - it is simply the expected number of characters that have evolved on *T* by 2 edge changes and that also satisfy the three conditions (C-i)–(C-iii) above. By Equation (27), this can be approximated by 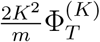. Note that this quantity is independent of λ (provided this is sufficiently small that the above approximations are reasonable), and *n* also is not involved in the first term in the product.

Note that one cannot interpret *mp_T_* as the expected number of false splits in a maximum compatibility tree involving the *K* perfectly evolved characters and the additional characters with 2 changes. This is because some of these latter characters may be incompatible with each other (even though they are compatible with the *K* perfectly evolved characters).

#### The case where ℓ > 1 binary characters evolve with 2 edge changes on T

So far we have considered the impact of a single binary character that has evolved on *T* with 2 edge changes. What happens if there is more than one such character? In particular, when will two such characters (which induce false splits) be compatible with each other? The following result provides a concise characterisation. This result is also relevant to case for more than two such characters, since a collection of binary characters is compatible if and only if every pair of characters is compatible.

##### Proposition 3

Suppose that *f* and *f*′ are two binary characters that have evolved on *T*, each with with 2 edge changes (say *e*_1_, *e*_2_ for *f* and 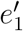 and 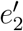 for *f*′). Let *P* (respectively, *P*′) denote the set of edges in the path in *T* consisting of *e*_1_, *e*_2_ (respectively 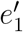 and 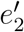) and the edges on the path between them. Then *f* and *f*′ are compatible if and only if either one of the following two conditions holds:

(*c*_1_) *P* ⊆ *P*′ or *P*′ ⊆ *P*.
(*c*_2_) *e* does not lie in *P*′ and *e*′ does not lie in *P*.

*Proof*: The proof involves a case analysis. In particular, we show that:

1. if (*c*_1_) holds then *f* and *f*′ are compatible;
2. if (*c*_1_) fails but (*c*_2_) holds, then *f* and *f*′ are compatible; and
3. if (*c*_1_) and (*c*_2_) both fail to hold, then *f* and *f*′ are incompatible.

To simplify notation we first introduce some conventions. We will let *σ* (respectively *σ*′) denote the split of [*n*] induced by (reversing state) changes on *e*_1_ and *e*_2_ (respectively on 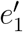 and 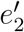). For subsets *V*_1_, …, *V_r_* of [*n*] we write *V*_1_ … *V_r_* | – as shorthand for the split (*V*_1_ ∪ … ∪ *V_r_*)|([*n*] – (*V*_1_ ∪ … *V_r_*)), and if a subtree within *T* has leaf set *A* ⊂ [*n*] we will denote this subtree by writing *t*(*A*). We will also assume that *e*_1_, *e*_2_, 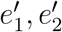 are four distinct edges (we deal with the case where this fails at the end).

Case (1): Suppose that (*c*_1_) holds. Without loss of generality we may assume that *P* ⊆ *P*′ and that the order of the path from 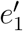 to 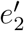 passes through *e*_1_ and *e*_2_ in this order. Thus we can represent *T* as in Fig. 2(i) where *A, B, C, D, E* denote the (unions of the) leaf sets of the corresponding subtrees determined by this arrangement of the four edges.

Thus *σ* is the split *ABDE*|*C*, and *σ*′ is the split *AE*|*BCD*. Since the second half of he first split (namely *C*) is a subset of the second half of the second split (*BCD*) these two splits are compatible. Note that this argument also holds if one or both of the following identities applies: 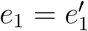 or 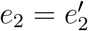.

Case (2): Now suppose that (*c*_1_) fails but (*c*_2_) holds. In this case, by considering the path between *e*_1_ and *e*_2_ we can represent *T* as in Fig. 2(ii) and 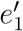 and 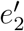 either both lie the subtree *t*(*A*) or *t*(*C*) or else 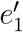 lies in *t*(*B_i_*) and 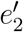 lies in *t*(*B_j_*) for some *i*, *j* (we allow *i* = *j*)

First suppose that 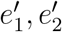 both lie within *t*(*A*). Then *σ*′ = *A*′| – for a subset of *A*′ of *A* and since *σ* = *AC*|*B* these two splits are compatible. A similar argument holds if 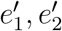 both lie within *t*(*C*). Alternatively, suppose that 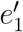 is an edge in *t*(*B_i_*) and 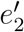 is an edge of *t*(*B_j_*) (we allow *i* = *j*). Then *σ*′ = *B*′| – for a subset of *B*, and since *σ* = *AC*|– these two splits are also compatible (since 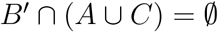).

Case (3): Finally suppose that Case (*c*_1_) and Case (*c*_2_) both fail. Without loss of generality we may assume that *e*_1_ lies within *P*′. In this case we can represent tree *T* as shown in Fig. 2(iii). Thus *σ*′ = (*A* ∪ *D*)| –.

By the assumption that Case (*c*_1_) and Case (*c*_2_) both fail, it follows that *e*_2_ lies in one of the following trees *t*(*A*), *t*(*D*), *t*(*B_i_*), or *t*(*C_j_*), for some *i*, *j*. By symmetry, there are just two sub-cases to consider: (i) *e*_2_ lies in *t*(*A*) or (ii) *e*_2_ lies in *t*(*B_i_*). In subcase (i) *σ* = *CDA*′|– where *A*′ is a proper subset of *A*, and *C* is the union of the leaf sets in *C*_1_, … *C_s_*. Thus *σ* = *CDA*′|– is incompatible with *σ*′ = *AD*|– since neither of the sets on the left of the split contains the other.

In subcase (ii), *σ* = *B*′*CD*|– for a proper subset *B*′ of *B* (the strict containment is because *e*_1_ and *e*_2_ are not adjacent) and so again *σ* and *σ*′ are incompatible.

Finally, we assumed that *e*_1_, *e*_2_, 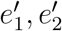 are four distinct edges. Otherwise (since *e*_1_ ≠ *e*_2_ and 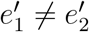) we either have: 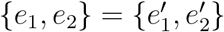, in which case condition (*c*_1_) holds, and the two characters *f* and *f*′ induce the same split and so are compatible, or we may assume without loss of generality that 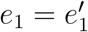 and 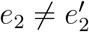. A similar (though simpler) case analysis to the above leads to the same conclusions as before.

**Fig. 2.**
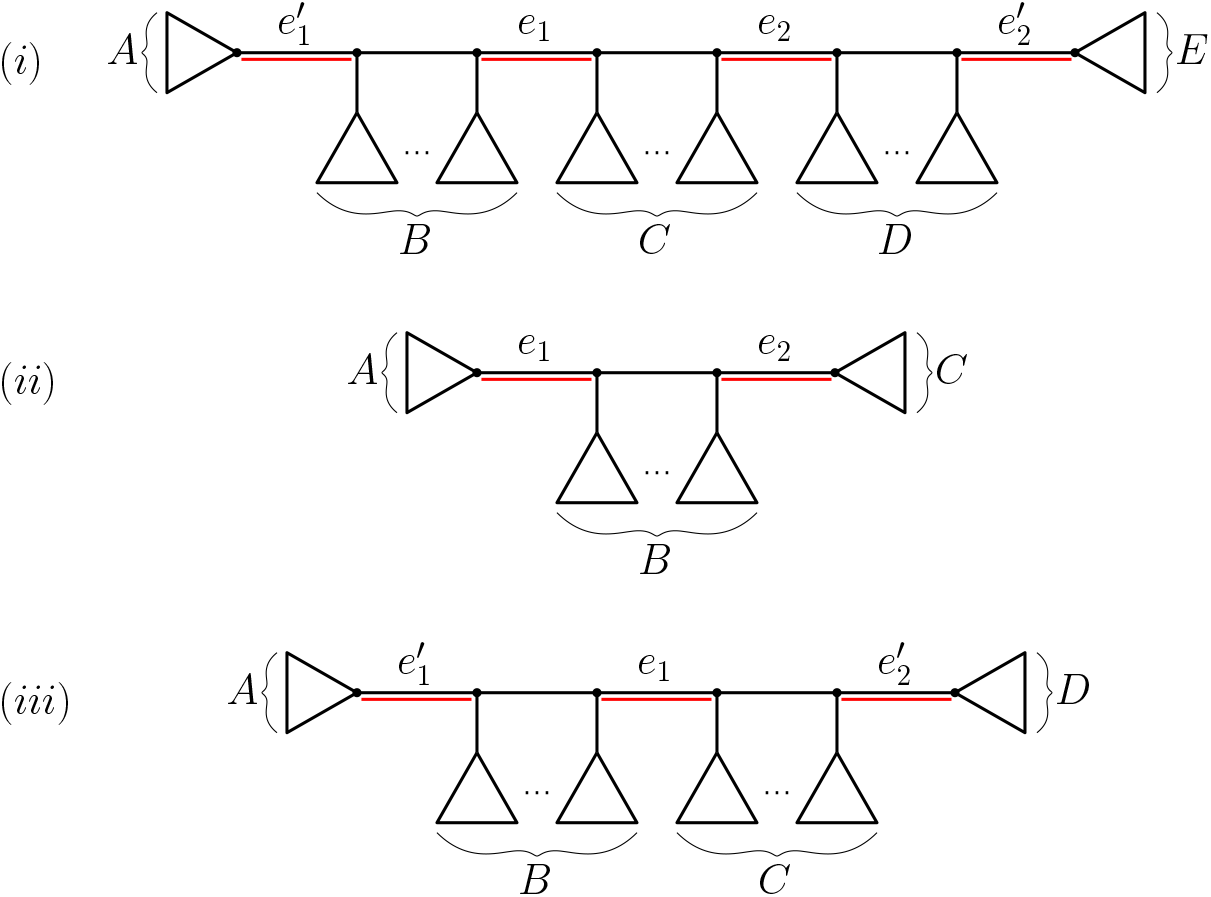
The cases for the proof of Proposition 3

##### Remark

A consequence of Proposition 3 and our earlier results is that if *n* is large in comparison to *k* then a small number (ℓ) of 2-state characters evolved randomly on *T* are likely to be (i) compatible with each other, (ii) compatible with the *k* perfectly evolved characters, and (iii) not be splits of *T*, and thus show up as false splits on the reconstructed (max compatabiilty or max parsimony) tree. To see this, observe first that the probability that (*c*_2_) occurs tends to 1 as *n* → ∞ for fixed *k* (this holds for any choice of *T*, or a randomly sampled tree from the PDA or YH distribution^†^. Second, a collection of binary characters is compatible if and only if each pair of them is compatible, and so if ℓ and *k* are fixed (or grow sufficiently slowly with *n*, depending on the tree shape) the probability that all ℓ false splits of *T* are present in the MP or MC tree (along with the splits corresponding to the *k* perfectly compatible characters of *T*) tends to 1 as *n* → ∞.

##### Estimating the false positive rate

Given a binary phylogenetic tree *T*, and *m* characters evolved randomly on *T* by the model described earlier, the *false positive rate* (*FP_T_*) is the expected value of the proportion of non-trivial splits in the reconstructed tree (using MC, say) that are not in *T* (here we assume that if the reconstructed tree is a star, this proportion (which is technically 0/0) is zero). Recall that *ξ* is the expected number of state changes in the tree *T* per character, under the model described earlier. *FP_T_* is a function of the three parameters *T* (specifically, its shape and number of leaves), *m* and λ (equivalently, *FP_T_* is a function of *T*, *m* and *ξ*).

In general, it is mathematically complicated to describe *FP_T_* in terms of these parameters. However, when the number of leaves in a tree grows faster than the number of perfectly compatible characters, it is possible to state a limit result, in order to provide an approximation to *FP_T_* for large trees.

In the following theorem we consider the following setting:

I. *mξ* = Θ(*n^β^*) for some 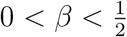, and
II. *mξ*^2^ = *O*(1),

where *O*(1) refers to dependence on *n* (thus *mξ*^2^ is not growing with *n*). Note that Condition (I) implies that the number of perfectly-evolved characters grows with the number of leaves, but at a rate that is slower than linearly. When 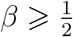, Condition (I) also provides a positive probability that more than one perfectly-evolved character will give rise to the same non-trivial split. Conditions (I) and (II) imply that *ξ* decreases as *n* increases.

We will show that in this setting, the false positive rate is (asymptotically) of the form 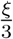 times a function Ω that involves *T* (via its shape), *m* and *ξ*. To describe this result, we need to define this function Ω. Let

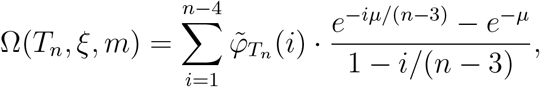

where

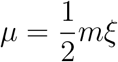

and where 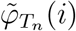 is given in Eqn. (2). For example, for any caterpillar tree, we have 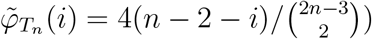.

Notice that Ω(*T_n_*, *ξ*, *m*) depends on *T_n_* only via the coefficients 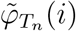, and this dependence in linear. Thus, if 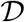 is a distribution on trees (e.g. the PDA or YH) then the expected value of Ω(*T_n_*, *ξ*, *m*) is given by:

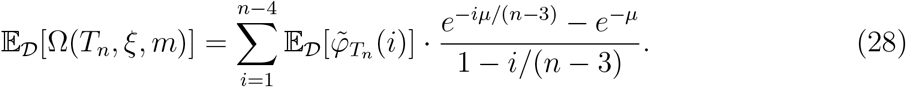

For the PDA distribution, the term 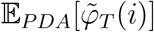 has an explicit exact value, given by Eqn. (9).

###### Theorem 3

i. 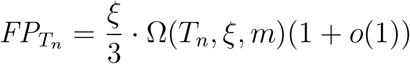

where *o*(1) is a term that tends to 0 as *n* grows.
ii. If *T_n_* is sampled from a distribution 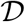 (e.g. PDA, YH) then the expected value of *FP_T_n__*, denoted 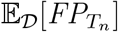, satisfies

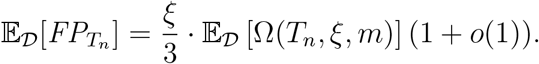
iii. If *T_n_* is a caterpillar tree, then *FP_T_n__* = 4(1 + *o*(1))/(3*m*).

*Proof: Part (i):* For convenience, we will write *T* in place of *T_n_* The number *X*_1_ of perfectly evolved characters on *T* has a binomial distribution (*m* trials, with the probability of success of 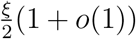 (since the proportion of interior edge in *T* is 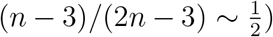 and so has expected value 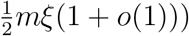. As before, let *K* denote the number of distinct non-trivial splits of *T_n_* that correspond to (one or more) of these perfectly evolved characters. By Condition (I) it follows that each perfectly evolved character is (with probability → 1) generated at most once, and so *K* is approximated by *X*_1_, which in turn is approximated (for large *n*) by a Poisson distribution with mean 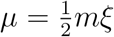.

Consider now the splits that arise from 2-edge-change characters on *T*. Denote the number of false splits by *X*_2_, and the number of true splits by 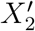. The expected value of 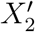 is of order *mξ*^2^/*n* and so it converges to zero as *n* grows, by Condition (II). By Proposition 3 and Condition (II), the probability that every pair of the *X*_2_ false splits is compatible (with each other) tends to 1 as *n* grows.

Next, consider splits (true or false) that arise from characters involving 3 or more changes on different edges. If *X*_3_ denotes the number of such characters, then the expected value of *X*_3_ is bounded above by a constant (independent of *n*) times *mξ*^3^, and by Conditions (I) and (II), the ratio of this to *K* is of order *ξ*^2^ (independent of *n*).

Thus,

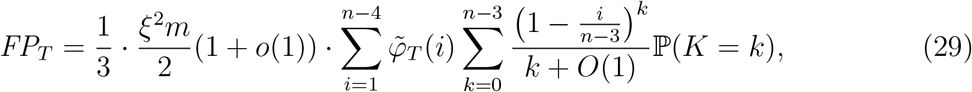

where *K* has a Poisson distribution with expected value 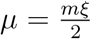, and *O*(1) refers to a term that is bounded in *n* (by Condition (II)) and accounts for any non-trivial splits induced by characters that are not perfectly evolved and in the reconstructed tree (as well as for splits from perfectly evolved characters that are ‘lost’ in a strict consensus by being the only such split in a path between the two edges of a 2-change character), while *o*(1) refers to a term the tends to zero as *n* grows due to characters that involve 3 or more edge changes).

Thus, 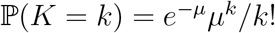, and so, under Conditions (I) and (II) we have:

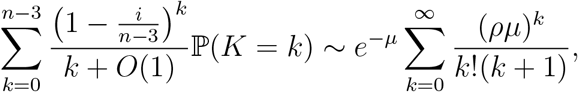

where 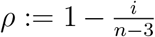. We now apply the identity: 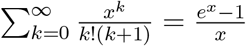, with *x* = *ρμ* to obtain:

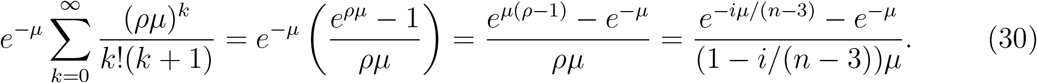

Since *μ* = *mξ*/2, notice that we can write the term 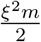 in Eqn. (29) as *ξμ* and so, from Eqns. (29) and (30), we have:

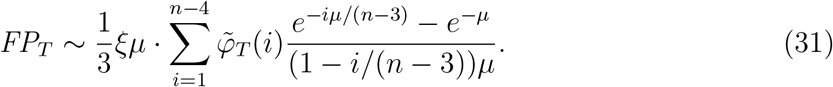

Finally, canceling the term *μ* in the numerator and denominator on the right-hand-side of Eqn. (31) gives the expression in Part (i).

*Part (ii):* This follows directly from Part (i) by linearity of expectation.

*Part (iii):* When *T_n_* is a caterpillar tree we have:

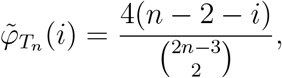

for 1 ⩽ *i* ⩽ *n* – 3. We can rewrite this as: 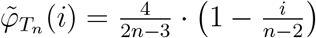, and substituting this into Part (i) we obtain:

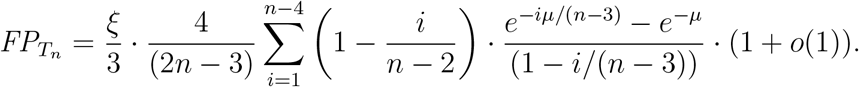

As *n* → ∞ we have the asymptotic identity:

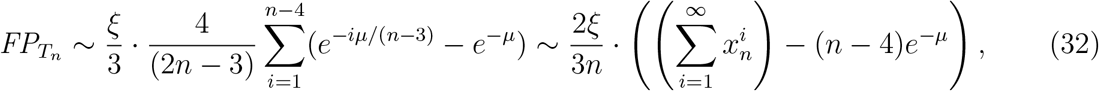

where 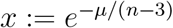.

Now, *x* converges to 1 from below as *n* grows (under Conditions (I) and (II)), and so applying the identity 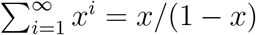 (for 0 < *x* < 1), together with the identity 1 – *x* ~ *μ*/(*n* – 3) ~ *μ*/*n* = *mξ*/2*n* (since 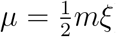) gives:

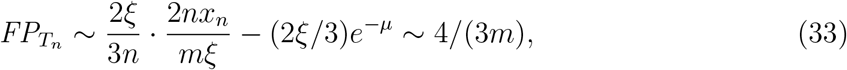

noting that the term (2*ξ*/3)*e*^−*μ*^ is asymptotically negligible relative to the term it follows in Eqn. (33) under Conditions (I) and (II). This completes the proof.

## Appendix 1: Links between ML and MP

Given *T* ∈ *B*(*n*) and a data set *D* consisting of a sequence of *m* characters. Let *n_i_* be the number of these characters that have parsimony score *i* on *T*, for *i* ⩾ 0 (thus *n*_0_ is the number of constant-state characters present in the data, *n*_1_ is the number of characters present that could have perfectly evolved on *T* etc). We will assume that (i) each character has a unique most-parsimonious representation on *T* and (ii) no edge is involved in a state change for more than one character (under a most-parsimonious representation on *T*).

These conditions are reasonable under conditions (1) and (2) described earlier for large *n*, where *n*_0_ >> *n*_1_ >> *n*_2_… and where *m* grows more slowly than *n* so changes on edges are likely to be ‘well-spaced’. We consider a likelihood setting where an edge counted by *n_i_* has probability *p_i_* and an edge not used in any most parsimonious reconstruction has probability *v*.

Note that

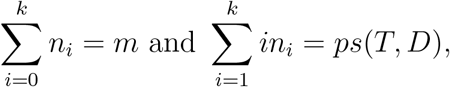

where *ps*(*T, D*) is the parsimony score of the tree *T* for the data. We will assume that *ps*(*T, D*) << *m*.

Now, we can write the likelihood function for *T* as follows:

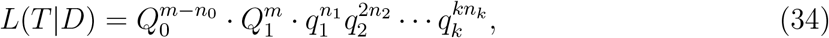

where 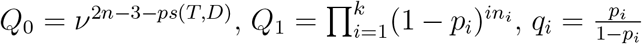. Clearly, *L*(*T*|*D*) is maximised by setting *v* = 1 (regardless of the other parameters). Moreover, the log likelihood critical values for *p_i_* obtained by solving the equations:

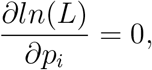

gives 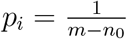 for all *i* ⩾ 1; in particular, the optimal *p_i_* values are all equal, and applying Eqn. (34) shows that *L*(*T*|*D*) is then a monotone decreasing function of *ps*(*T, D*). In this case, as noted in (2) (p. 353), the MP tree(s) maximize this optimal likelihood score.

## Appendix 2

The probability a false split does not appear

Now consider any given binary tree *T*, a character *f* that has evolved on *T* by 2 edge changes on *e*_1_, *e*_2_, together with *k* perfectly evolved characters on *T* (with all the characters independently evolved under the Jukes-Cantor model with *p_e_* constant across the edges of *T*).

Let 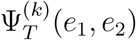 be the probability that *f* satisfies conditions (C-i) and (C-ii) above (i.e. it is binary, and corresponds to a split that is not in *T*), and this split does **not** occur in either the MP (or MC) tree for the data that consists of *f* together with *k* perfectly evolved characters. In the case of ties (in constructing the MP or MC tree) these are broken uniformly.

### Proposition 4

Then:

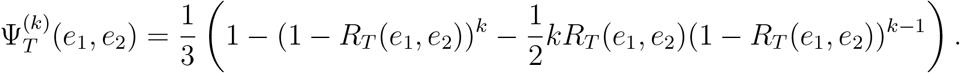

*Proof*: Let 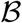 be a binomial random variable consisting of *k* trials, with the probability of success on each trial of 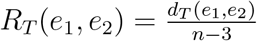 (as defined earlier). Then 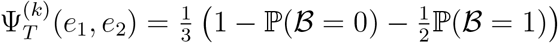. The factor of 1/2 is to allow for the breaking of a tie when 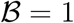 between the two MC (or MP) trees.

Observe that if *e*_1_ and *e*_2_ are adjacent, then *R_T_*(*e*_1_, *e*_2_) = 0 and so 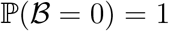 and thus 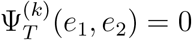.

Next, let 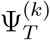 be the expected value of 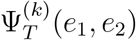 across all pairs of edges {*e*_1_, *e*_2_}. Thus, 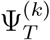 is a natural quantity to compare with the earlier 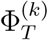. We then have:

### Corollary 2

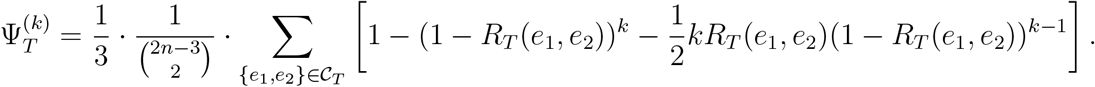

### Remark

Notice that 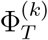 is a monotone decreasing function of *k* (and thus 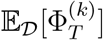 is also). Also, since we can write 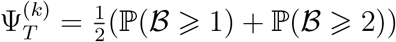 it follows that 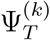 is a monotone increasing function of *k* (and so 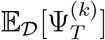 is also).

Notice also that

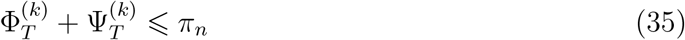

and

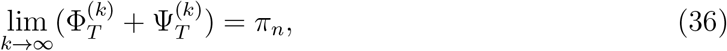

where 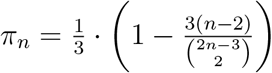 is the conditional probability that a character *f* is binary and that the split induced by this binary character is incompatible with *T* given that *f* has evolved on *T* with 2 edge changes. To see this, observe that there are 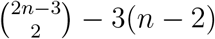 pairs of non-adjacent edges in *T*, all 2 edge change characters have the same probability. Eqns. (35) and (36) also apply if Φ and Ψ are replaced by their expected value over a tree distribution 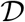.

---

**Table 1.** Full Set of Parameters of 11 Virus Phylogenies Sampled^1^

| Abbrev | $n$ | $m$ | $\xi$ | $\alpha_e$ | Res | $r_{AC}$ | $r_{AG}$ | $r_{AT}$ | $r_{CG}$ | $r_{CT}$ | $r_{GT}$ | $f_A$ | $f_C$ | $f_G$ | $f_T$ | $\alpha_{ASR}$ | $p_{INV}$ |
| --- | --- | --- | --- | --- | --- | --- | --- | --- | --- | --- | --- | --- | --- | --- | --- | --- | --- |
| DENV | 1197 | 10264 | 6.68 | 0.08 | 0.88 | 1.15 | 7.83 | 1.55 | 0.80 | 23.74 | 1 | 0.32 | 0.21 | 0.26 | 0.21 | 0.44 | 0.29 |
| DENV-1 | 1067 | 10264 | 2.88 | 0.31 | 0.82 | 1.03 | 7.02 | 1.31 | 0.74 | 18.86 | 1 | 0.32 | 0.21 | 0.26 | 0.21 | 0.38 | 0.45 |
| EBOV | 1610 | 18164 | 0.15 | 0.51 | 0.36 | 0.99 | 7.16 | 0.62 | 0.33 | 8.48 | 1 | 0.32 | 0.21 | 0.20 | 0.27 | 0.29 | 0.66 |
| EV-D68 | 824 | 7293 | 2.83 | 0.29 | 0.80 | 1.44 | 31.73 | 1.88 | 1.71 | 46.62 | 1 | 0.32 | 0.20 | 0.21 | 0.27 | 0.26 | 0.53 |
| HIV | 189 | 1038 | 0.18 | 0.45 | 0.22 | 1.51 | 4.24 | 0.66 | 0.20 | 3.39 | 1 | 0.39 | 0.16 | 0.21 | 0.23 | 0.02 | 0.79 |
| MeV | 109 | 15782 | 0.44 | 0.23 | 0.70 | 1.39 | 9.47 | 0.62 | 0.40 | 13.81 | 1 | 0.29 | 0.24 | 0.23 | 0.23 | 0.44 | 0.55 |
| MuV | 458 | 15154 | 0.07 | 0.23 | 0.30 | 0.63 | 3.04 | 0.16 | 0.14 | 3.62 | 1 | 0.31 | 0.22 | 0.20 | 0.27 | 0.07 | 0.75 |
| RSV | 997 | 14986 | 0.88 | 0.29 | 0.61 | 1.40 | 8.61 | 1.35 | 0.25 | 15.78 | 1 | 0.39 | 0.18 | 0.16 | 0.28 | 0.42 | 0.52 |
| SARS-CoV-2 | 583 | 29668 | 0.02 | 0.66 | 0.23 | 0.44 | 1.26 | 0.18 | 0.39 | 3.81 | 1 | 0.30 | 0.18 | 0.20 | 0.32 | 0.02 | 0.90 |
| WNV | 2512 | 10395 | 2.03 | 0.57 | 0.60 | 1.08 | 5.22 | 1.14 | 0.54 | 16.05 | 1 | 0.27 | 0.22 | 0.29 | 0.22 | 0.37 | 0.52 |
| ZIKV | 543 | 10320 | 0.45 | 0.33 | 0.55 | 0.99 | 4.02 | 1.19 | 0.74 | 10.76 | 1 | 0.27 | 0.22 | 0.29 | 0.21 | 0.66 | 0.45 |

<sup>1</sup> $n$ =number of leaves; $m$ =number of sites; $\xi$ =tree length; Res=Fractional resolution on tree; $r_{XY}$ =rate parameter of the GTR matrix from base $X$ to $Y$ ; $f_X$ =frequency of base $X$ ; $\alpha_e$ =edge rate variation gamma shape parameter; $\alpha_{ASR}$ =across-site rate variation gamma shape parameter; $p_{INV}$ =fraction of invariant sites

**Fig. 3.**
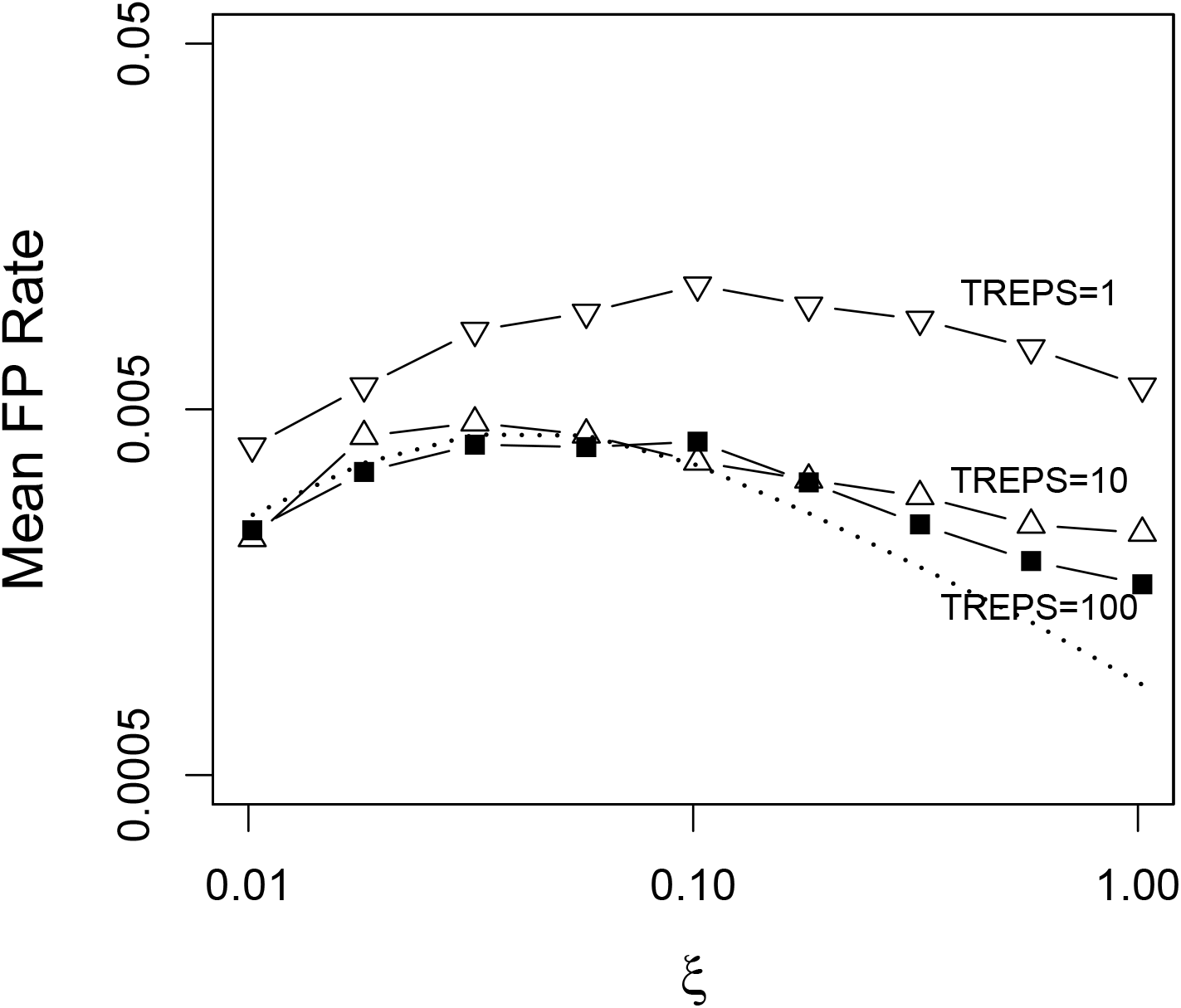
Mean expected false positive rate in the near-perfect zone of *ξ* ⩽ 1 for three different breadths of MP tree search algorithm for a PDA distribution of trees (1,10, and 100 random addition sub-replicates per simulation replicate; 1000 simulation replicates). Predicted values from Eqn. 2 (main text) is given by dashed line.

**Fig. 4.**
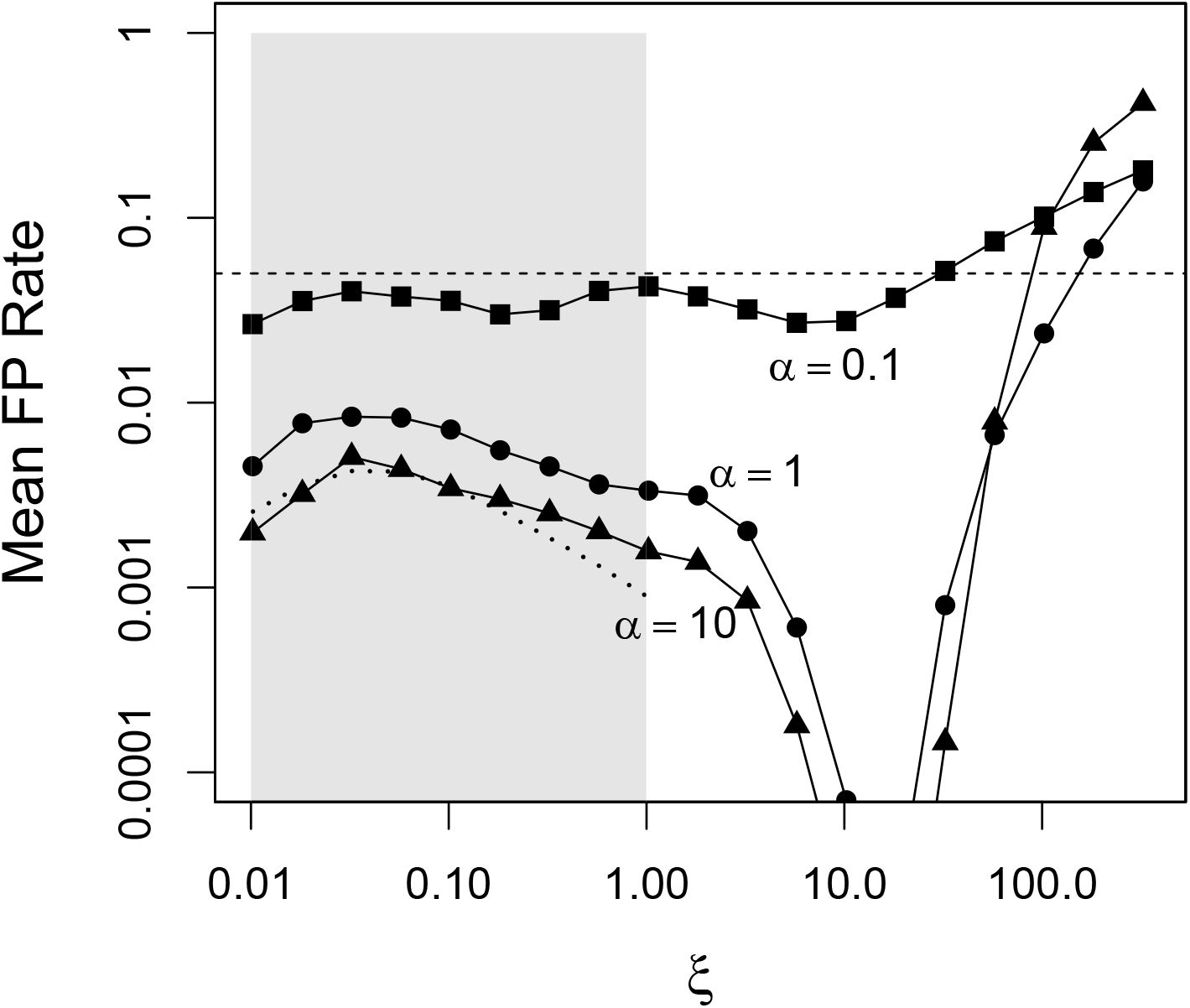
Effect of across-site-rate variation on expected false positive rate for MP inference for different values of the *α*_ASR_ shape parameter of the ASR gamma distribution. Smaller *α* values have higher rate variation. Edge length variation is assumed absent. Dashed curve is the prediction from Eqn. 2 (main text), in which both sources of variation are absent. Tree search algorithm has 100 subreplicates per simulation replicate; 1000 simulation replicates.

**Fig. 5.**
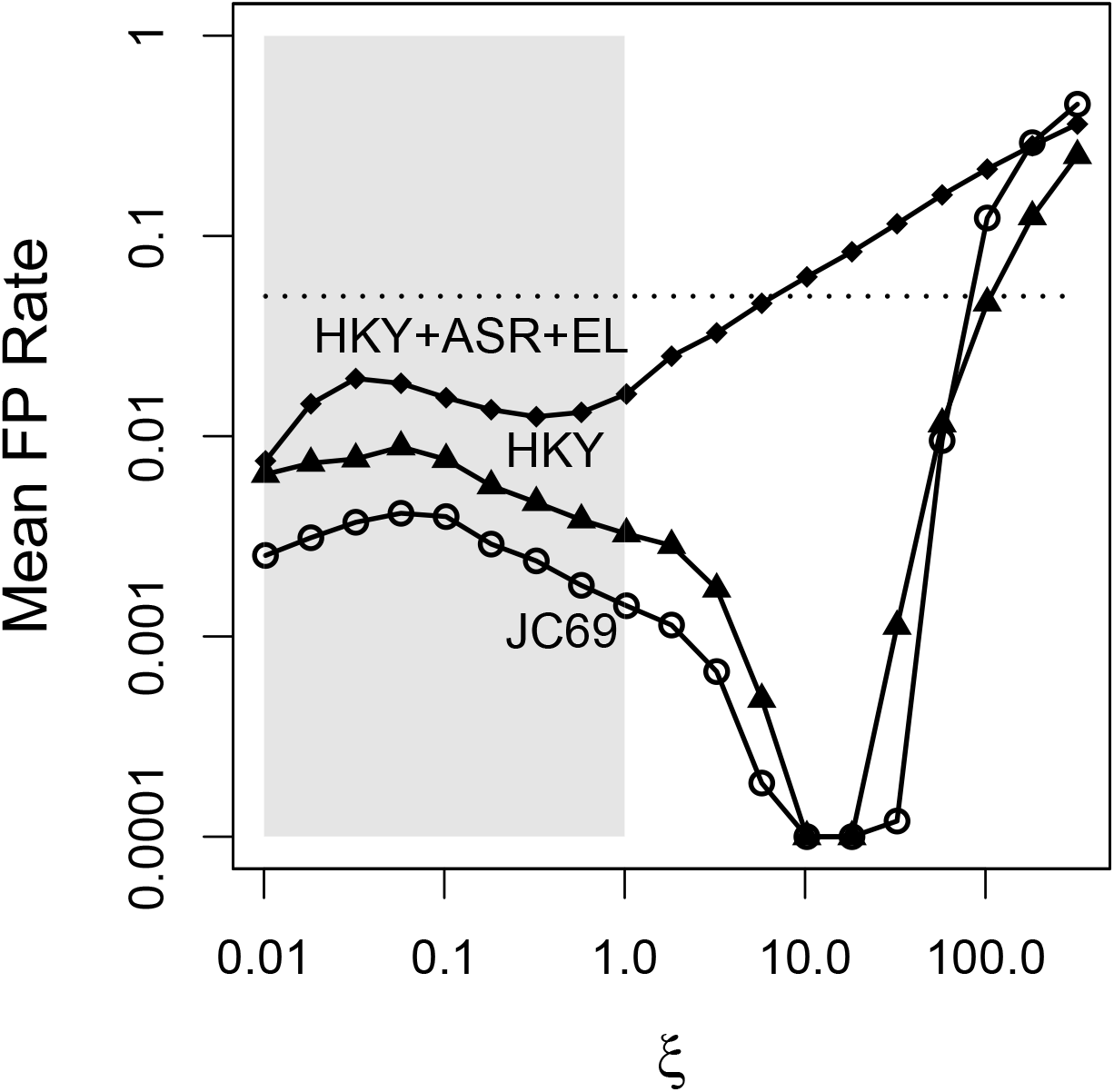
Effect of model complexity on expected false positive rates for MP. JC69 model (open circles) corresponds to near-perfect assumptions in shaded box (*ξ* ⩽ 1). HKY model (closed triangles) has a transition:transversion ratio of 5 and equal base frequencies. HKY+ASR+EL model (closed diamonds) has a transition:transversion ratio of 5 and also rate variation across sites (*α_ASR_* = 1) and edges ((*α*_e_ = 1)). Points are means of 1000 replicates × 100 sub-replicates.

**Fig. 6.**
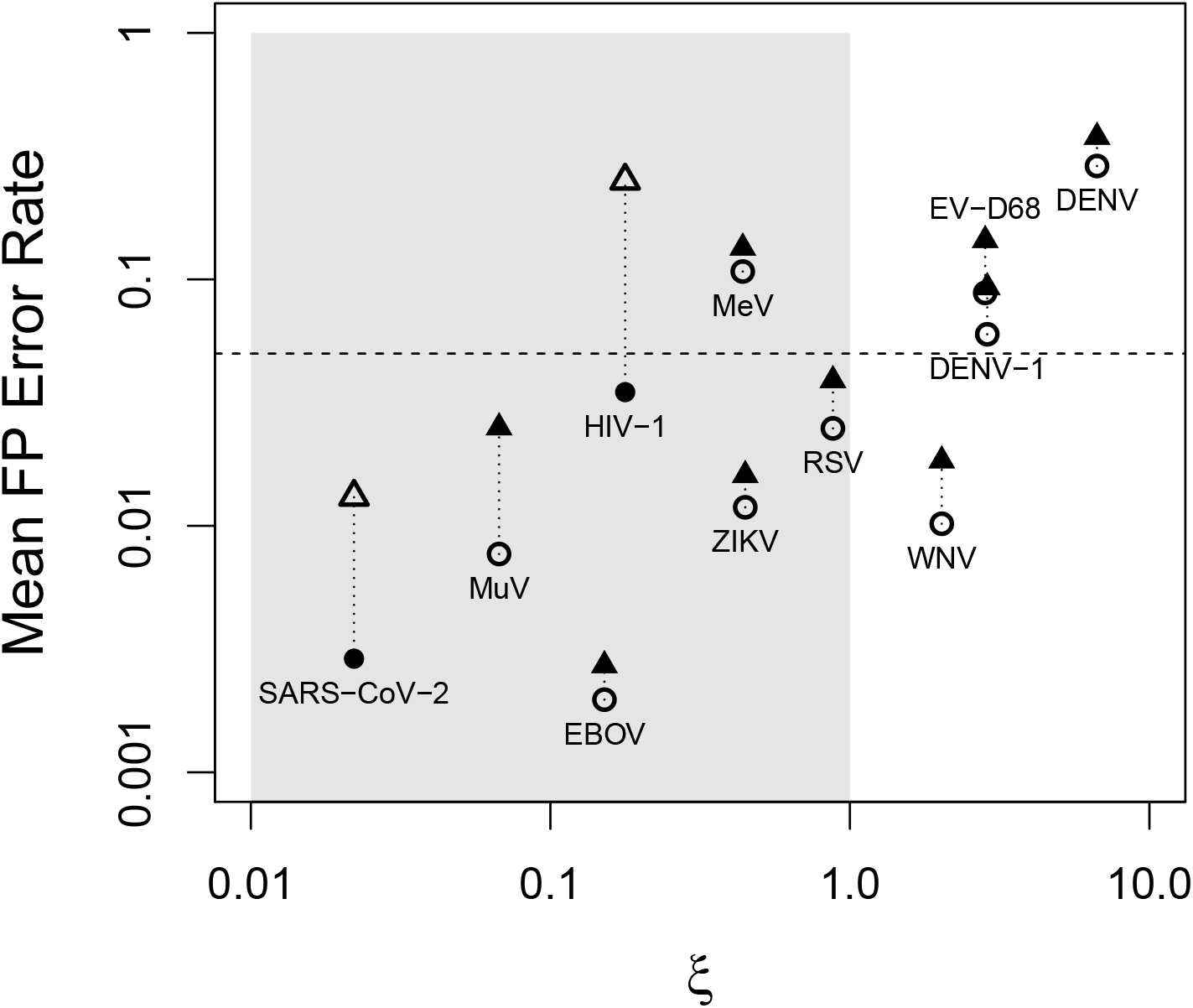
Mean false positive rates estimated for 11 viral phylogenetic data sets, as a function of *ξ* and other parameters estimated from the data. Simulations used parameters given in Table 1, and are for MP assuming ASR variation follows either an invariant sites model (circles) or a gamma distributed model (triangles), with higher likelihood model point shaded, all assuming a PDA distribution of trees (100 subreplicates per simulation replicate; 500 simulation replicates). Near-perfect zone of *ξ* ⩽ 1 is shaded. Horizontal dashed line indicates a 0.05 FP rate.

**Fig. 7.**
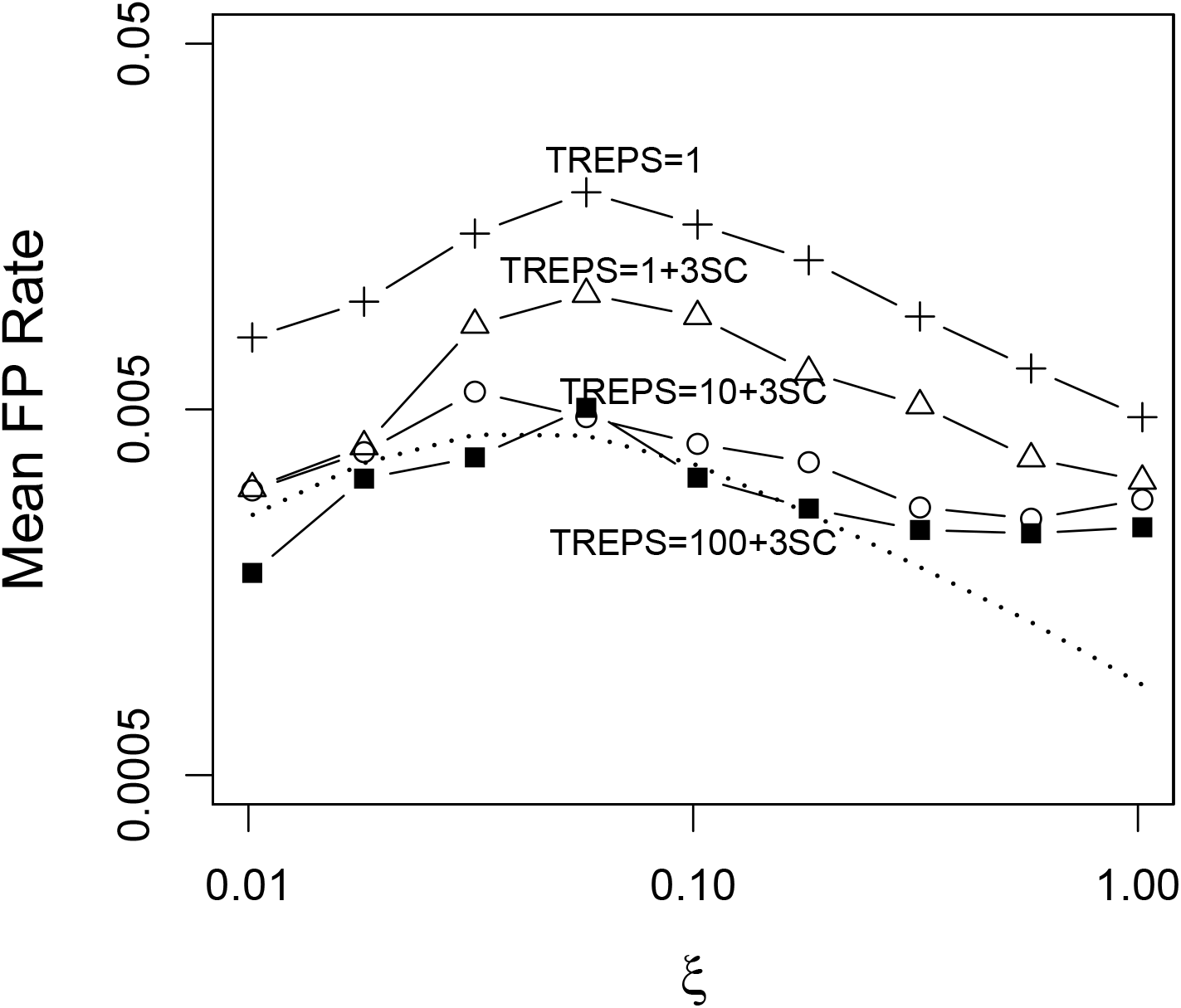
Mean false positive rate in the near-perfect zone of *ξ* ⩽ 1 for maximum likelihood searches in IQTree2 using different search breadths and corrections. Each point is the mean in 500 replicate simulations with a PDA distribution of trees. Open plusses: one subreplicate search per simulation replicate; open triangles: one subreplicate plus 3-state correction; open circles: 10 subreplicate searches plus 3-state correction and trees combined via strict consensus. closed squares:100 subreplicate searches plus 3-state correction and trees combined via strict consensus. The “3-state correction” collapses edges less than 1/*m* substitutions per site, prior to consensus (*m* = 1000 sites). For comparison to theory for MP, predicted values from Eqn. 2 are given by dashed line.

**Fig. 8.**
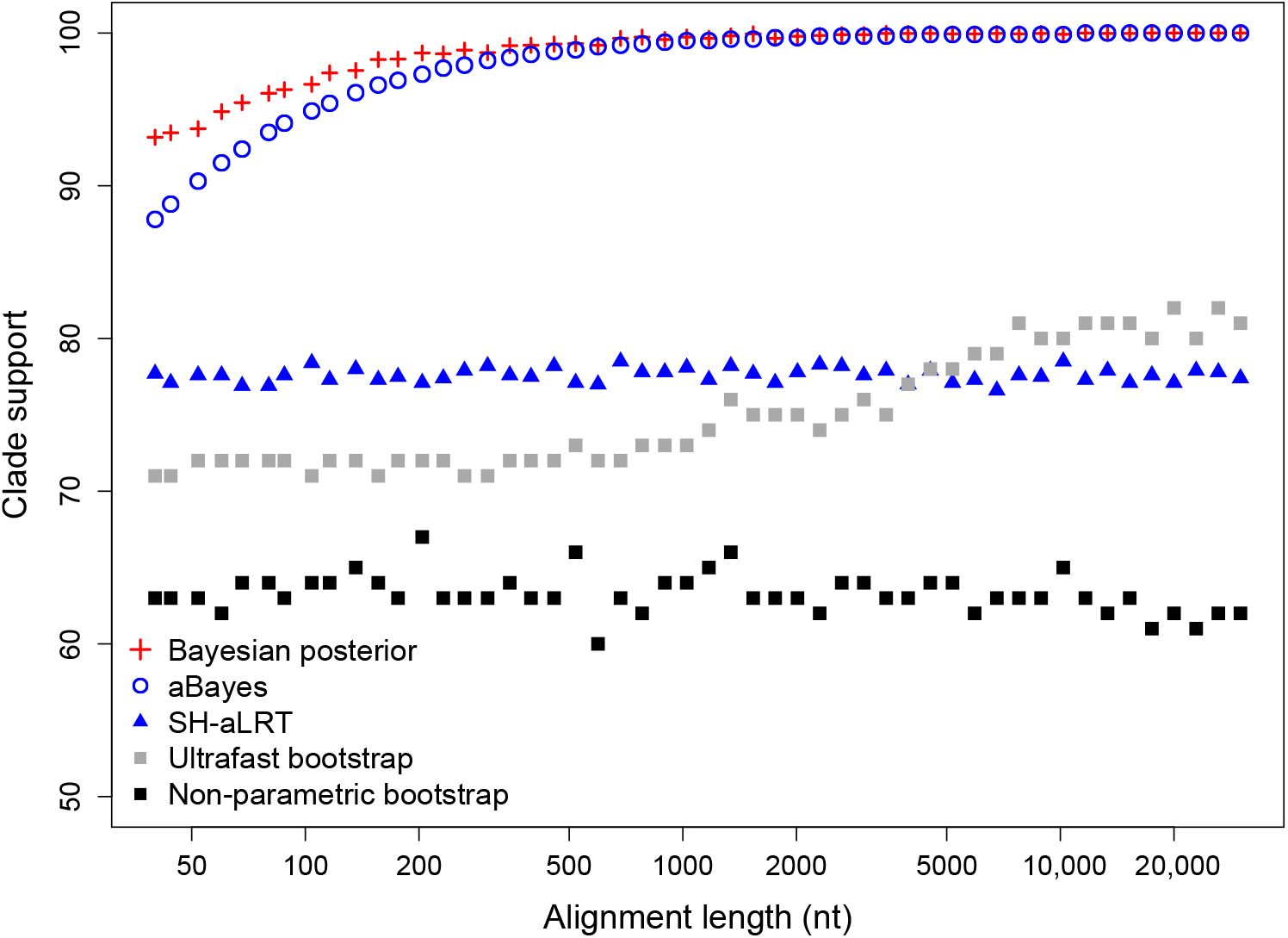
Five measures of clade support estimated on perfect four-taxon trees as alignment length varies. Each edge of the tree has exactly one site with a substitution on that edge; all other sites are constant. Support metrics using bootstrap resampling are denoted with solid shapes.

**Fig. 9.**
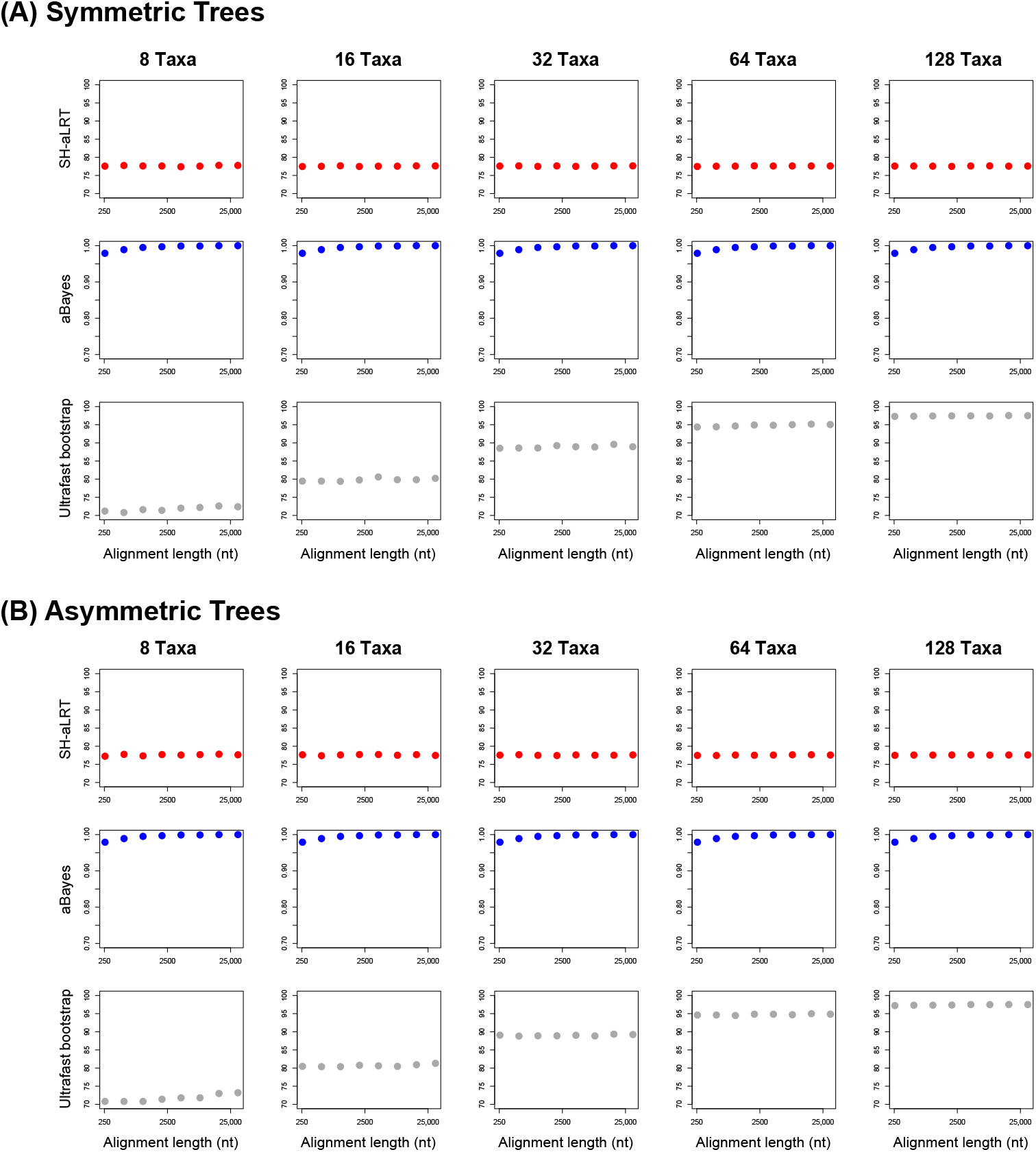
Three measures of clade support estimated on perfect trees of different sizes and alignment lengths. Each edge of the tree has exactly one site with a substitution on that edge; all other sites are constant. (A) Simulations on perfectly symmetric trees. (B) Simulations on perfectly asymmetric (caterpillar) trees.

